# Remdesivir and GS-441524 retain antiviral activity against Delta, Omicron, and other emergent SARS-CoV-2 variants

**DOI:** 10.1101/2022.02.09.479840

**Authors:** Jared Pitts, Jiani Li, Jason K. Perry, Venice Du Pont, Nicholas Riola, Lauren Rodriguez, Xianghan Lu, Chaitanya Kurhade, Xuping Xie, Gregory Camus, Savrina Manhas, Ross Martin, Pei-Yong Shi, Tomas Cihlar, Danielle P. Porter, Hongmei Mo, Evguenia Maiorova, John P. Bilello

## Abstract

Genetic variation of SARS-CoV-2 has resulted in the emergence and rapid spread of multiple variants throughout the pandemic, of which Omicron is currently the predominant variant circulating worldwide. SARS-CoV-2 variants of concern or interest (VOC/VOI) have evidence of increased viral transmission, disease severity, or decreased effectiveness of vaccines and neutralizing antibodies. Remdesivir (RDV, VEKLURY^®^) is a nucleoside analog prodrug and the first FDA-approved antiviral treatment of COVID-19. Here we present a comprehensive antiviral activity assessment of RDV and its parent nucleoside, GS-441524, against 10 current and former SARS-CoV-2 VOC/VOI clinical isolates by nucleoprotein ELISA and plaque reduction assay.

Delta and Omicron variants remained susceptible to RDV and GS-441524, with EC_50_ values 0.31 to 0.62-fold of those observed against the ancestral WA1 isolate. All other tested variants exhibited EC_50_ values ranging from 0.15 to 2.3-fold of the observed EC_50_ values against WA1. Analysis of nearly 6 million publicly available variant isolate sequences confirmed that Nsp12, the RNA-dependent RNA polymerase (RdRp) target of RDV and GS-441524, is highly conserved across variants with only 2 prevalent changes (P323L and G671S). Using recombinant viruses, both RDV and GS-441524 retained potency against all viruses containing frequent variant substitutions or their combination. Taken together, these results highlight the conserved nature of SARS-CoV-2 Nsp12 and provide evidence of sustained SARS-CoV-2 antiviral activity of RDV and GS-441524 across the tested variants. The observed pan-variant activity of RDV supports its continued use for the treatment of COVID-19 regardless of the SARS-CoV-2 variant.

## 1. Introduction

Since the emergence of severe acute respiratory syndrome coronavirus 2 (SARS-CoV-2) in late 2019, two lineages, numerous variants, and subvariants have been detected through genomic surveillance. As with other coronaviruses, SARS-CoV-2 variants emerge through inter- and intramolecular recombination and from heritable errors generated in the viral genome by its error-prone RNA-dependent RNA polymerase (RdRp) (1, 2). Retention of genetic changes in a viral genome may be linked to advantages in replication fitness and/or in overcoming selective pressures exerted by the host immune system, an antiviral, or a therapeutic neutralizing antibody. Naturally occurring SARS-CoV-2 variants can be categorized as variants of concern (VOC) or interest (VOI) by the World Health Organization (WHO) or Centers for Disease Control and Prevention (CDC) based on evidence of increased rates of transmission and disease severity, detection failures, or potential loss in susceptibility to current vaccines and neutralizing antibodies. The early ancestral A lineage isolates detected in Wuhan, China and Seattle, WA (WA1 strain) were rapidly replaced worldwide by the B lineage VOC Alpha in 2020 (3). Subsequently, multiple VOIs and dominant VOCs of the B lineage progressively emerged, with Delta and most recently Omicron completely replacing prior strains (4–6). The defining genetic changes that differentiate variants predominantly occur in the gene encoding the spike protein, which mediates virus binding, fusion, and entry. However, changes are also detected elsewhere in the viral genomes, resulting in infrequent amino acid substitutions in the Nsp5 3CL main protease (Mpro) and Nsp12 RdRp, the two targets of currently approved SARS-CoV-2 antivirals.

Remdesivir (RDV; VEKLURY^®^) (7) was the first antiviral approved for the treatment of patients hospitalized with COVID-19 based on evidence that RDV treatment significantly reduced recovery times in clinical trials (8–10). Further, in the PINETREE clinical trial, in which RDV was administered in an outpatient setting, RDV reduced COVID-19 related hospitalization and death by 87% (11). These results led to an expanded FDA approval of RDV for high-risk non-hospitalized individuals with COVID-19 symptoms (12).

RDV is a nucleotide mono-phosphoramidate prodrug of the parent nucleoside GS-441524 (13). Following IV administration, RDV is metabolized intracellularly to the active triphosphate metabolite (RDV-TP), effectively bypassing the rate-limiting first phosphorylation step of GS-441524. RDV-TP then competes efficiently with cellular ATP for incorporation into the nascent SARS-CoV-2 viral RNA, resulting in cessation of strand-synthesis by two separate mechanisms of action (14, 15). Prior to the emergence of SARS-CoV-2, RDV and its parent nucleoside GS-441524 were shown to inhibit multiple RNA viruses (16–18), including a broad spectrum of coronaviruses such as SARS-CoV, Middle Eastern respiratory syndrome coronavirus (MERS-CoV), mouse hepatitis virus (MHV), and other zoonotic coronaviruses (19–22). Additionally, potent antiviral activity of RDV was observed in primary lung cells *in vitro* and confirmed *in vivo* across multiple respiratory viruses including respiratory syncytial virus (RSV) (18), Nipah (23), SARS-CoV (20), and MERS-CoV (22).

The RdRp catalytic active site is nearly 100% conserved among coronaviruses, therefore the observed potency of RDV against other coronaviruses was anticipated to translate to SARS-CoV-2 antiviral activity (21). RDV and GS-441524 have both demonstrated potency against SARS-CoV-2, with *in vitro* cellular EC_50_ values ranging from 10 to 120 nM for RDV and 470 to 3600 nM for GS-441524 (13, 24–27). The *in vivo* efficacy of RDV has been demonstrated in SARS-CoV-2 challenge studies in mice and hamsters (13, 28, 29). Additionally, RDV efficacy was demonstrated in non-human primates following several different routes of administration including IV, SC, and inhalation (30–32).

The low sequence diversity and high genetic stability of the SARS-CoV-2 RNA replication complex, including the Nsp12 RdRp, observed over time indicates a minimal global risk of pre-existing resistance to RDV (33). However, the emergence of each new variant brings a risk of altered susceptibility to vaccine-induced immunity, therapeutic antibodies, or antivirals. In this report, we demonstrate that *in vitro* potencies of RDV and GS-441524 are preserved among the known prominent SARS-CoV-2 variants as well as against recombinant viruses harboring specific substitutions frequently observed in variant Nsp12.

## 2. Results

### Antiviral activity of RDV and GS-441524 against clinical isolates of SARS-CoV-2 variants

*In vitro* RDV antiviral activity was assessed against clinical isolates of an extensive panel of past and present SARS-CoV-2 VOC/VOIs (Supplemental Table 1). Antiviral activity was initially assessed utilizing a plaque reduction assay (PRA) from culture supernatants harvested at 48 hours post-infection (hpi) from A549-ACE2-TMPRSS2 cultures infected with variants at an MOI of 0.1. The average RDV EC_50_ value for the WA1 reference strain by PRA was 98 ± 48 nM, while variant EC_50_ values ranged from 15 to 154 nM, representing 0.15- to 1.6-fold changes relative to WA1 (Table 1 and Fig. 1A). These results indicate that RDV retains potent antiviral activity against all variants evaluated by PRA, including the Delta variant, which was ~3-fold more susceptible to RDV than the WA1 isolate.

**Figure 1.**
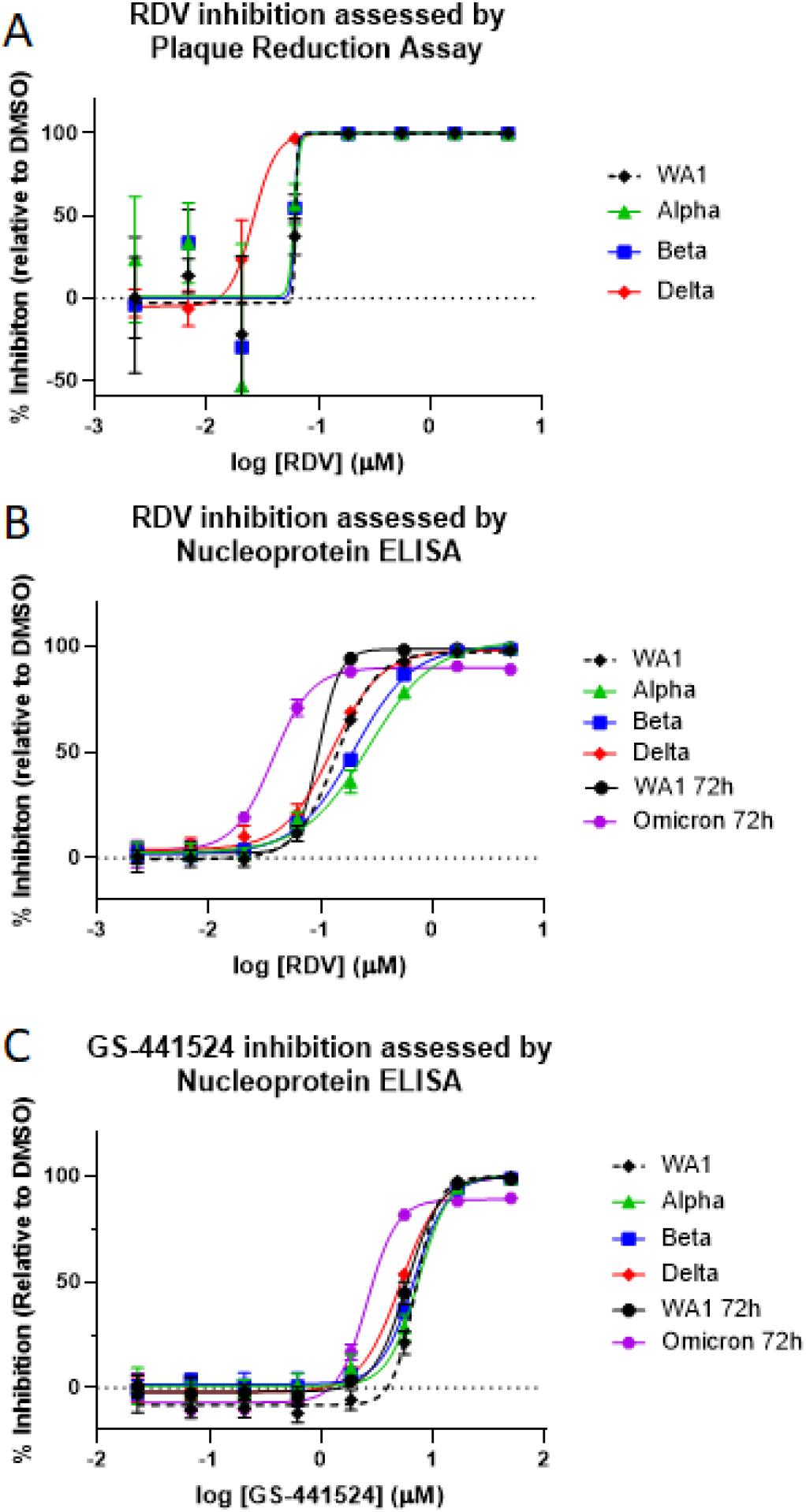
Representative dose-response curves of remdesivir and GS-441524 against SARS-CoV-2. **VOCs.** Dose-response curves of RDV (A-B) and GS-441524 (C) activity against the WA1 reference and SARS-CoV-2 VOCs in A549-ACE2-TMPRSS2 cells by plaque reduction assay (PRA) (A) or ELISA (B-C). In the PRA, infected cell supernatants were harvested at 48 hpi and analyzed by plaque assay on Vero-TMPRSS2 cells. For ELISA, infected cells were fixed at ~48 hpi (WA1, Alpha, Beta, Delta) or ~72 hpi (WA1 and Omicron) and processed. Data shown are means and standard deviations from representative experiments that were performed in biological quadruplicate (PRA) or triplicate (ELISA) at each compound concentration. Average calculated EC_50_ values and fold change from WA1 reference can be found in Table 1.

**Table 1.** Antiviral activity of RDV and GS-441524 against SARS-CoV-2 variants.

| Variant | RDV Plaque Reduction Assay (n = 2-4) |  | RDV Nucleoprotein ELISA (n = 4-16) |  | GS-441524 Nucleoprotein ELISA (n = 4-12) |  |
| --- | --- | --- | --- | --- | --- | --- |
|  | Mean EC<sub>50</sub> ± SD<sup>a</sup> (nM) | Fold Change from WA1<sup>b</sup> | Mean EC<sub>50</sub> ± SD<sup>a</sup> (nM) | Mean Fold Change ± SD from WA1<sup>c</sup> | Mean EC<sub>50</sub> ± SD<sup>a</sup> (nM) | Mean Fold Change ± SD from WA1<sup>c</sup> |
| WA1 | 98 ± 48 | 1.0 | 110 ± 42 | 1.0 | 5600 ± 4100 | 1.0 |
| Alpha | 94 ± 58 | 0.96 | 192 ± 51 | 1.58 ± 0.48 | 8790 ± 6600 | 1.22 ± 0.60 |
| Beta | 61 ± 9 | 0.63 | 141 ± 45 | 1.19 ± 0.47 | 7570 ± 4400 | 1.13 ± 0.56 |
| Gamma | 154 ± 226 | 1.6 | 97 ± 39 | 0.82 ± 0.42 | 5060 ± 2300 | 0.79 ± 0.37 |
| Delta | 31 ± 13 | 0.31 | 70 ± 40 | 0.59 ± 0.20 | 3260 ± 1300 | 0.62 ± 0.24 |
| Epsilon | 65 ± 32 | 0.66 | 210 ± 212 | 1.94 ± 1.18 | 4050 ± 1700 | 1.27 ± 0.53 |
| Zeta | 87 ± 44 | 0.89 | 151 ± 102 | 1.17 ± 0.40 | 3840 ± 1400 | 0.93 ± 0.11 |
| Iota | 59 ± 28 | 0.60 | 258 ± 195<sup>d</sup> | 2.33 ± 0.74 | 4710 ± 1600 | 1.43 ± 0.28 |
| Kappa | 15 ± 6 | 0.15 | 77 ± 50 | 0.63 ± 0.19 | 2100 ± 930 | 0.53 ± 0.07 |
| Lambda | 94 ± 55 | 0.96 | 175 ± 138 | 1.37 ± 0.48 | 3890 ± 1600 | 0.97 ± 0.10 |
| WA1 (72 hpi) | - | - | 97 ± 15 | 1.0 | 6240 ± 1300 | 1.0 |
| Omicron (72 hpi) | - | - | 44 ± 16<sup>e</sup> | 0.45 ± 0.13 | 3330 ± 1400<sup>e</sup> | 0.57 ± 0.29 |

<sup>a</sup> Values are the mean ± standard deviation (SD) of the results of independent experiments (number of replicate experiments shown).
<sup>b</sup> Fold change calculated from the mean values = [Variant mean EC<sub>50</sub>]/[WA1 mean EC<sub>50</sub>]
<sup>c</sup> A fold change was calculated for each experiment and a mean fold change ± SD was calculated with these values.
<sup>d</sup> Statistically significant increase (p=0.015) in EC<sub>50</sub> value of Iota in the RDV ELISA compared to WA1 reference at 48 hpi by one-way ANOVA with Bonferroni correction for multiple comparisons. All other results for variants at 48 hpi are not statistically different from matching WA1 reference.
<sup>e</sup> Statistically significant decrease (p≤0.0001) in EC<sub>50</sub> value of Omicron RDV and GS-441524 ELISA compared to WA1 reference at 72 hpi by one-way ANOVA with Bonferroni correction for multiple comparisons.

To assess the antiviral effect directly in infected cultures, antiviral testing based on the nucleoprotein (N) enzyme-linked immunoassay (ELISA) was developed and conducted in addition to PRA. The EC_50_ of RDV against the WA1 reference strain by ELISA was 110 ± 42 nM (Table 1 and Fig. 1B), indicating that antiviral potency of RDV is consistent between PRA and ELISA. Variants tested in ELISA had average RDV EC_50_ values ranging from 70 to 258 nM with observed fold EC_50_ changes from 0.59 to 2.33 (Table 1 and Fig. 1B). Due to the low signal observed with the Omicron variant at the standard 48 h timepoint, likely stemming from the reduced *in vitro* replication efficiency of Omicron (34), the assay was extended to 72 h and compared with WA1 assessed at the same timepoint. The WA1 reference isolate at 72 hpi had an average RDV EC_50_ of 97 ± 15 nM. Thus, delaying the readout to 72 hpi had no effect on the observed potency of RDV. The Omicron variant was significantly more susceptible to RDV with an average EC_50_ value of 44 ± 16 nM (p≤0.0001), a 0.45-fold change compared to WA1.

The potency of GS-441524, the parent nucleoside of RDV, was also unchanged against all variants and the WA1 reference isolate, as measured by ELISA at 48 hpi. The EC_50_ of GS- 441524 at this timepoint against WA1 was 5600 ± 4100 nM and ranged from 2100 to 8790 nM (0.53- to 1.43-fold change from WA1) against the collection of SARS-CoV-2 variants (Table 1, Fig. 1C). In agreement with the findings for RDV, GS-441524 was also found to be significantly (p ≤0.0001) more potent against Omicron than WA1 at 72 hpi, with EC_50_ values of 3330 ± 1400 compared to 6240 ± 1300 nM for WA1.

### Nsp12 sequence changes in SARS-CoV-2 variants

To assess the genetic variation of Nsp12 in the 11 current or previously classified VOC/VOIs (Omicron, Delta, Alpha, Beta, Gamma, Epsilon, Zeta, Iota, Kappa, Lambda, and Mu), a total of 5,842,948 SARS-CoV-2 variant sequences from the GISAID (Global Initiative on Sharing Avian Influenza Data) database were evaluated. The highest proportion of analyzed sequences were Delta variants (4,059,836; 69.5%), followed by Alpha variants (1,158,351; 19.8%) and Omicron variants (392,056; 6.7%); the other 8 variants made up the remaining 4.0% (Supplemental Table 2). We further assessed the genetic variation in spike in comparison to Nsp12 across the variants.

The number of amino acid substitutions from WA1 viral isolate sequence in Nsp12 and spike was calculated for each of the 11 variants. Overall, 1 to 6 amino acid substitutions were observed across the different variants, with a frequency of ≥1% of sequences over the 932 amino acid positions in Nsp12 compared with a range of 7 to 45 substitutions over the 1274 amino acid positions in spike (Supplemental Fig. 1, Supplemental Tables 2 and 3). The most prevalent Nsp12 substitution relative to the consensus ancestral sequence, P323L, was observed with frequency >99% and a lineage defining Nsp12 substitution for all 11 analyzed variants. The Delta variant contained one additional lineage-defining amino acid change in Nsp12, G671S, which was observed in 97.8% of Delta isolates. No other substitutions were found with a frequency of ≥50% in any of the variants.

Given the recent emergence and high prevalence of the Omicron variant, amino acid substitutions in Nsp12 of Omicron variant were further investigated with a more sensitive frequency cutoff of 0.5%. Among the 6 substitutions (Table 2), P323L and F694Y were the most frequently observed, in 99.5% and 2.0% of Omicron sequences, respectively, while all 4 remaining substitutions had frequencies of ≤1%. In the initial Omicron wave (December 13, 2021), F694Y was highly prevalent (41.1% of worldwide isolates and 94.1% of the UK isolates) in sequences submitted to the GISAID database; however, as the Omicron variant continued to spread, the G694Y substitution rapidly declined in frequency with only 2.00% of deposited sequences harboring the substitution as of January 18, 2022 (Supplemental Fig. 2). Interestingly, F694Y is not unique to the Omicron variant as it was also found in 4.9% of Delta variant sequences (Supplemental Table 2).

**Table 2.**
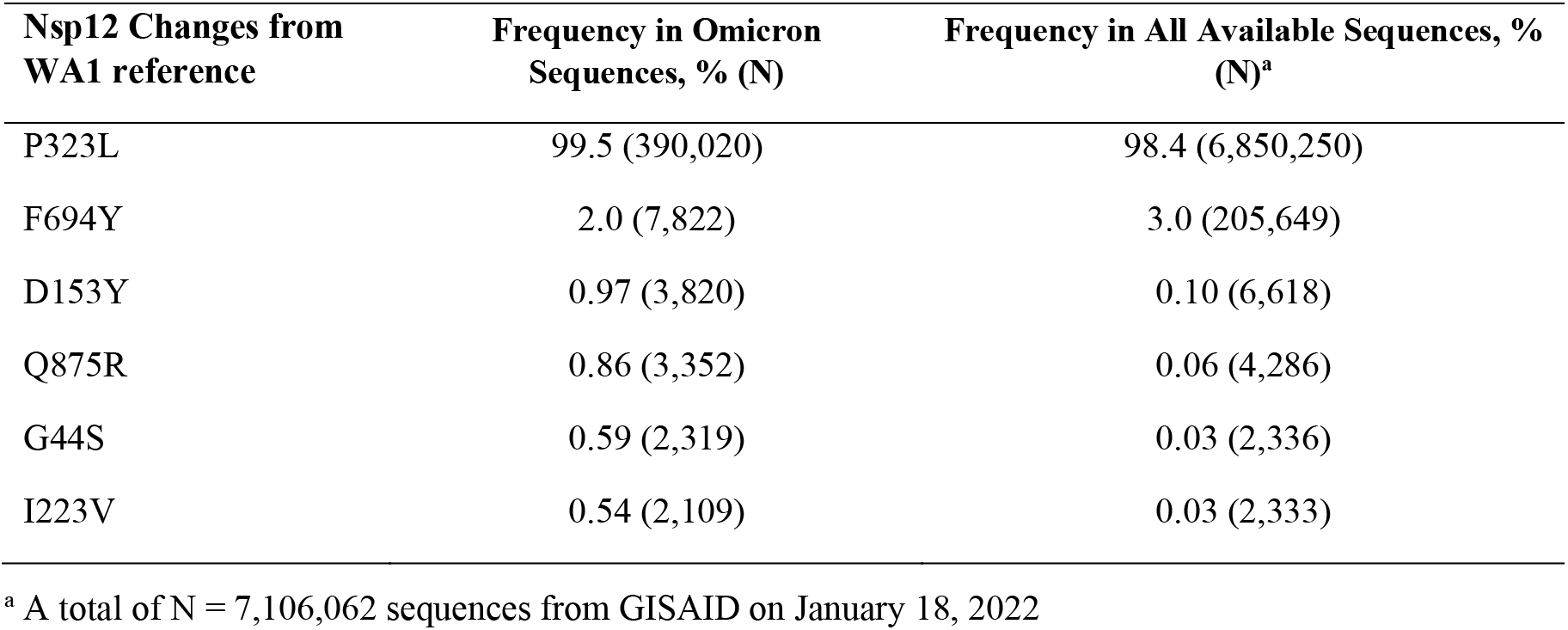
Amino acid substitutions in Nsp12 Omicron sequences with frequency ≥0.5%.

| Nsp12 Changes from<br>WA1 reference | Frequency in Omicron<br>Sequences, % (N) | Frequency in All Available Sequences, %<br>(N)<sup>a</sup> |
| --- | --- | --- |
| P323L | 99.5 (390,020) | 98.4 (6,850,250) |
| F694Y | 2.0 (7,822) | 3.0 (205,649) |
| D153Y | 0.97 (3,820) | 0.10 (6,618) |
| Q875R | 0.86 (3,352) | 0.06 (4,286) |
| G44S | 0.59 (2,319) | 0.03 (2,336) |
| I223V | 0.54 (2,109) | 0.03 (2,333) |

<sup>a</sup> A total of N = 7,106,062 sequences from GISAID on January 18, 2022

Most notably, Nsp12 substitutions previously identified to reduce *in vitro* susceptibility to RDV (19, 35), F480L, V557L, and E802D, were rarely found in our evaluation. Of the 5,842,948 variant sequences evaluated, F480L, V557L, and E802D were only observed in 16 (0.0002%), 24 (0.0004%), and 102 (0.002%) sequences, respectively. Further, only 49 (0.0008%) variant sequences had any alteration at the residue involved in RDV induced delayed chain termination, S861 (14).

### Structural analysis of Nsp12 substitutions observed in variants

RDV acts by incorporating its triphosphate metabolite (RDV-TP) into the viral RNA and subsequently causing clashes with the Nsp12 protein at multiple location, compromising further synthesis (14, 15). Analogs of RDV that produce the same RDV-TP active metabolite, such as GS-441524, exert their inhibitory activity via the same mechanism of action. At present, a structure of the pre-incorporated state of RDV-TP in the RdRp active site is still unavailable. We built a model, described previously (14, 15, 36), based on an existing structure of the polymerase complex (Nsp12/(Nsp8)2/Nsp7/(Nsp13)2) with primer and template RNA (PDB: 6XEZ) (37). Using this model, we assessed the potential impact of each Nsp12 amino acid substitution identified in the analyzed variants on the affinity of RDV-TP for the RdRp active site.

As seen in Fig. 2, the two most common amino acid substitutions, P323L, seen in all variants, and G671S, observed in Delta, are 28.6 Å and 24.9 Å, respectively, from the pre-incorporated RDV-TP (measured from the amino acid Cα to RDV-TP’s C1’). Of all the low-frequency substitutions identified, only F694Y, found in 2-5% of Omicron and Delta isolates, is in close proximity to the RdRp active site. Measured to be 12.2 Å from the RDV-TP, the residue is not in direct contact with the inhibitor but is close enough to have an indirect conformational effect. However, an evaluation of its impact on RDV-TP binding affinity using a molecular mechanics generalized Born surface area (MM-GBSA) approach resulted in no meaningful difference, likely because of the relatively conservative change from phenylalanine to tyrosine (38).

**Figure 2.**
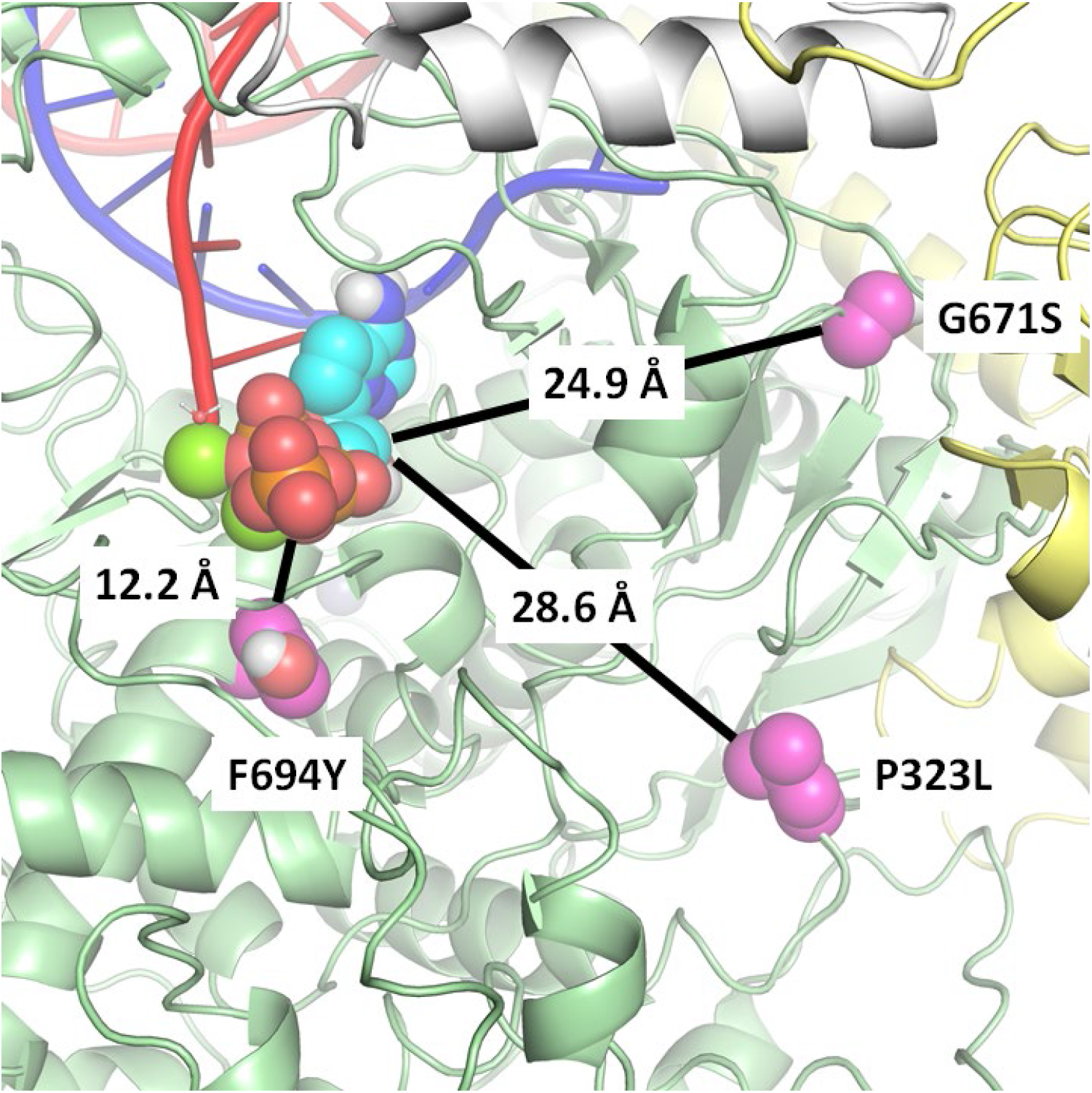
Structural model of Nsp12 highlighting key amino acid substitutions in relation to the active site. Pre-incorporated remdesivir triphosphate (RDV-TP) was modeled into the cryo-EM structure of the polymerase complex (6XEZ) (37). The prevalent amino acid substitution P323L, seen in all variants, is measured to be 28.6 Å from RDV-TP (P323 Cα - RDV-TP C1’), whereas G671S, seen in the Delta variant, is 24.9 Å. Of all the amino acid substitutions reported here, F694Y, seen at low frequency in Delta and Omicron, comes closest to the active site, at 12.2 Å. A computational analysis suggests that the substitutions have no meaningful impact on RDV-TP binding affinity.

Most low-frequency amino acid substitutions in Omicron and other variants occur on the surface of Nsp12, away from the polymerase active site (Supplemental Figs. 3 and 4). While the dynamics of incorporation and RDV-TP inhibition are complex events, this structural analysis suggests little reason to expect a significant impact on the efficacy of RDV and GS-441524.

### Potency against recombinant SARS-CoV-2 expressing the prevalent variant amino acid substitutions

The Omicron clinical isolate evaluated did not contain the Nsp12 F694Y substitution found at high frequency in early UK Omicron isolates (Supplemental Fig. 2). Due to the initial prevalence in Omicron variants and proximity to the RdRp active site, we sought to assess RDV and GS- 441524 activity against recombinant Omicron (rOmicron) viruses with and without the Nsp12 F694Y substitution. By ELISA, the RDV EC_50_ values were 46 ± 6 nM and 34 ± 3 nM (Table 3 and Fig. 3) against rOmicron and rOmicron F694Y, respectively. Similarly, potency was preserved for GS-441524 against both recombinant viruses, with EC_50_ values of 2600 ± 100 nM (rOmicron) and 2200 ± 300 nM (rOmicronF694Y). RDV and GS-441524 were similarly potent against the two recombinant Omicron viruses and an Omicron clinical isolate run in parallel, with all three viruses showing increased susceptibility to both RDV and GS-441524 compared to the WA1 isolate by ELISA (Table 3 and Fig. 3).

**Figure 3.**
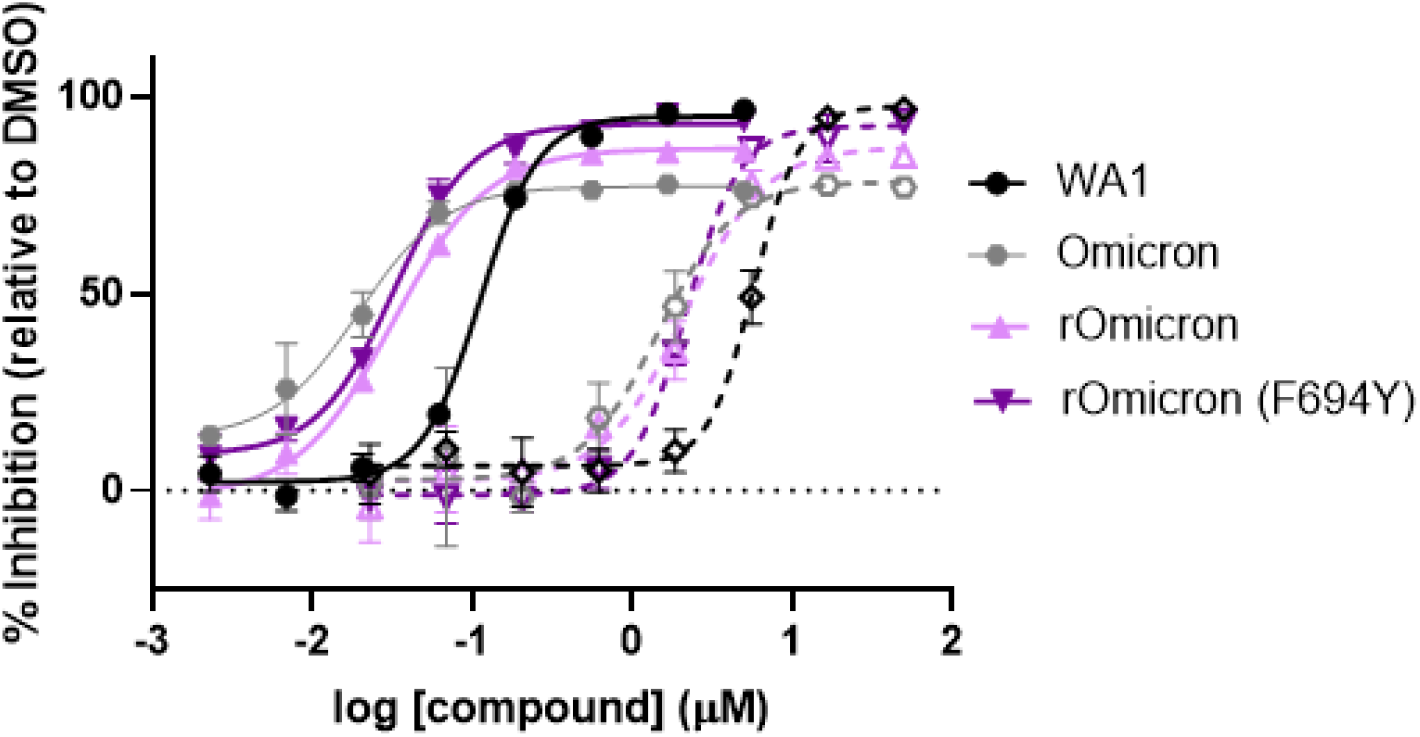
Recombinant Omicron viruses retain susceptibility to RDV and GS-441524. ELISA doseresponse curves of RDV (solid lines and filled points) and GS-441524 (dashed lines and outlined points) activity against recombinant Omicron viruses with (dark purple) or without (light purple) the F694Y substitution compared with WA1(black), and Omicron (grey) clinical isolates run in parallel. Data shown are means and standard deviations from a representative 72-hpi nucleoprotein ELISA experiment that was performed with biological triplicates at each compound concentration. Average calculated EC_50_ values are in Table 3.

**Table 3.** RDV and GS-441524 potencies against recombinant Omicron SARS-CoV-2 viruses.

| Virus | Mean EC<sub>50</sub> ± SD<sup>a</sup> (nM) |  |
| --- | --- | --- |
|  | RDV (n=2-3) | GS-441524 (n=2-3) |
| WA1 | 100 ± 15 | 5200 ± 700 |
| Omicron | 54 ± 21<sup>b</sup> | 4000 ± 1500<sup>b</sup> |
| rOmicron | 46 ± 6<sup>b</sup> | 2600 ± 100<sup>b</sup> |
| rOmicron (F694Y) | 34 ± 3<sup>b</sup> | 2200 ± 300<sup>b</sup> |

<sup>a</sup> Values are the mean ± standard deviation (SD) of the results of independent experiments (number of replicate experiments shown).
<sup>b</sup> Statistically significant decrease ( $p \leq 0.005$ ) in EC<sub>50</sub> value of Omicron and rOmicron viruses from the WA1 reference at 72 hpi by one-way ANOVA with Bonferroni correction for multiple comparisons. No statistical differences were observed between any of the Omicron and rOmicron viruses.

We next sought to evaluate the *in vitro* potency of RDV and GS-441524 against other Nsp12 amino acid substitutions alone or in combination that were identified at high frequency (>15%) in any specific variant. Recombinant SARS-CoV-2 WA1 viruses containing a Nano luciferase (Nluc) transgene and the wild-type or mutated nsp12 sequences were rescued and tested for RDV and GS-441524 susceptibility. Monitoring Nluc signal from infected cells at 48 hpi, we observed an RDV EC_50_ value of 80 ± 21 nM for WA1 recombinant virus, while viruses containing either the P323L substitution alone or the P323L/G671S double substitution found in Delta variant had RDV EC_50_ values of 71± 26 nM and 104 ± 20 nM, respectively, resulting in a 1.2-fold change or less relative to WA1 (Table 4, Fig. 4A). Modification of nsp12 with a sequence encoding G671S alone failed to rescue infectious virus after several independent attempts, a finding which complements prior evidence suggesting Nsp12 P323L conveys a growth advantage (39). Recombinant viruses containing either P323L, F694Y, or the P323L/F694Y double substitution in a WA1 Firefly luciferase (Fluc) recombinant virus background were similarly susceptible to RDV (EC_50_ values within 1.2-fold of WA1) (Table 4, Fig. 4B). GS-441524 antiviral potency was also maintained against recombinant SARS-CoV-2 viruses harboring the Nsp12 P323L, P323L/G671S, and P323L/F694Y substitutions, with fold changes of 0.73 to1.83 relative to WA1 (Table 4). Collectively, these data confirm that antiviral potencies of RDV and GS-441524 remain unchanged against viruses harboring the prevalent Nsp12 substitutions currently identified in isolates of SARS-CoV-2 variants.

**Figure 4.**
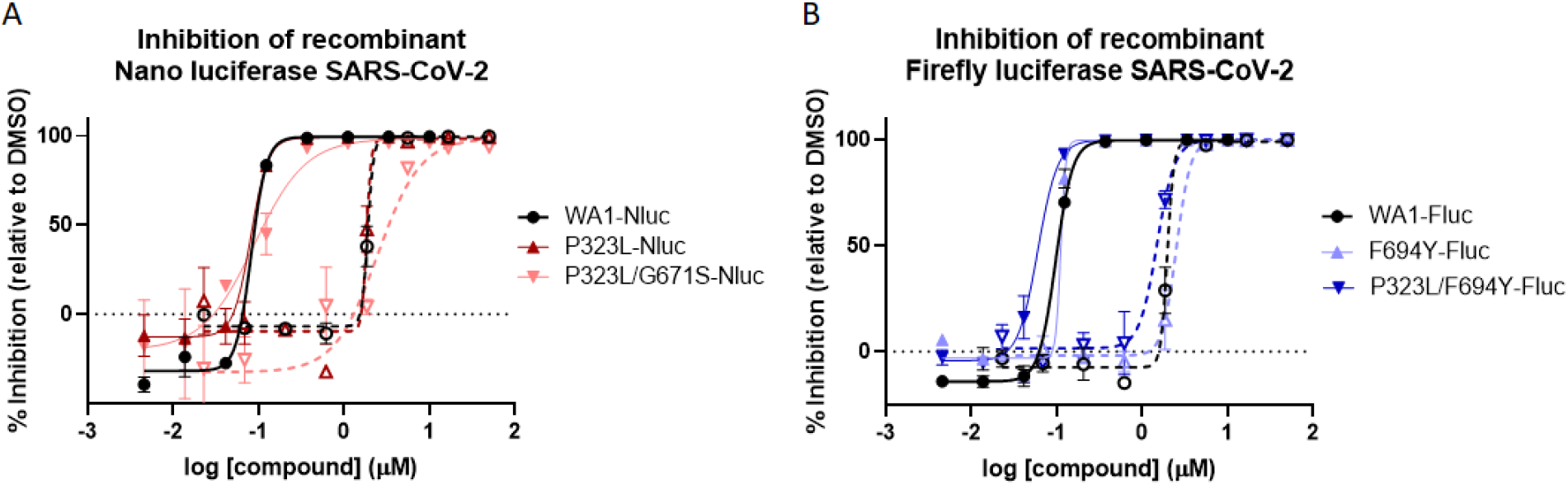
Prevalent Nsp12 substitutions in Delta and Omicron retain susceptibility to RDV and GS- 441524. Dose response curves of RDV (solid lines and filled points) and GS-441524 (dashed lines and outlined points) activity against recombinant viruses with/without prevalent Nsp12 substitutions containing a Nano luciferase (Nluc) (A) or Firefly luciferase (Fluc) (B) transgene. Data shown are means and standard deviations from a representative experiment that was performed in biological duplicates at each compound concentration. Average calculated EC_50_ values and fold change from recombinant WA1 references are in Table 4.

**Table 4.** RDV and GS-441524 potencies against recombinant SARS-CoV-2 harboring prevalent Nsp12 substitutions.

| Recombinant Virus | RDV (n=2-6) |  | GS-441524 (n=2) |  |
| --- | --- | --- | --- | --- |
|  | Mean EC<sub>50</sub> ± SD<sup>a</sup><br>(nM) | Mean Fold<br>Change ± SD from<br>WA1<sup>b</sup> | Mean EC<sub>50</sub> ± SD<sup>a</sup><br>(nM) | Mean Fold<br>Change ± SD from<br>WA1<sup>b</sup> |
| WA1-Nluc | 80 ± 21 | 1.0 | 1880 ± 40 | 1.0 |
| P323L-Nluc | 71 ± 26 | 0.95 ± 0.42 | 1580 ± 370 | 0.84 ± 0.21 |
| P323L/G671S-Nluc | 104 ± 20 | 1.22 ± 0.31 | 3450 ± 1400 | 1.83 ± 0.71 |
| WA1-FLuc | 99 ± 2 | 1.0 | 2230 ± 380 | 1.0 |
| F694Y-FLuc | 112 ± 2 | 1.14 ± .01 | 2250 ± 380 | 1.04 ± 0.34 |
| P323L/F694Y-FLuc | 63 ± 4 | 0.63 ± 0.03 | 1620 ± 180 | 0.73 ± 0.04 |

<sup>a</sup> Values are the mean ± standard deviation (SD) of the results of independent experiments (number of replicate experiments shown).
<sup>b</sup> A fold change was calculated for each experiment and a mean fold change ± SD was calculated with these values (number of independent replicate experiments shown).

### 3. Discussion and Conclusions

Over the past 2 years of the SARS-CoV-2 pandemic, the rapid evolution of the virus has led to the emergence of multiple viral variants. Since the beginning of pandemic, WHO declared 5 of these variants as VOCs that could be associated with more severe disease and/or increased rate of transmission. While the SARS-CoV-2 antiviral activity of RDV has previously been well characterized both *in vitro* and *in vivo* (13, 24, 26, 29–31), most studies have been conducted using the ancestral WA1 isolate. Here, we sought to fully characterize the antiviral potency of RDV and its parent nucleoside GS-441524 against a panel of the most significant SARS-CoV-2 variants including all the major VOCs. Utilizing PRA and ELISA assays in parallel, we observed a general agreement in potency between assays with most RDV EC_50_ values observed near 100 nM, indicating N-protein levels correlated with released infectious virus. Findings from both assays revealed all variants to have RDV EC_50_ values within 2.4-fold of WA1. The Iota variant was the only variant with >2-fold change in potency for RDV compared with WA1 by ELISA. However, the EC_50_ against Iota observed by PRA and for GS-441524 ELISA were similar to WA1 indicating that the Iota variant remains susceptible to RDV and GS-441524. Importantly, both Delta and Omicron variants, the two most recent strains in predominant circulation with increased severity and elevated transmission, respectively, are highly susceptible to both RDV and GS-441524. Interestingly, Omicron is significantly more susceptible to RDV and GS- 441524, though the reasons for this are not understood, as there are no substitutions in Nsp12 that would predict increased potency.

The potency observed for the GS-441524 parent nucleoside against all variants was 20-75 times lower than for RDV, consistent with previous findings in A549 cells (13). As observed with RDV, GS-441524 maintained potency against all clinical isolates of variants tested, with a maximum fold change of 1.4 compared with WA1. Although the active triphosphate for GS- 441524 and RDV are identical, it was important to confirm pan-variant GS-441524 potency because orally bioavailable prodrug options for delivery of GS-441524 are under exploration (40, 41).

The sequence analysis presented here and by others (22, 42) have found the nsp12 gene, encoding the RNA dependent RNA polymerase, of variants to be remarkably stable over the last 2 years. Only two substitutions, P323L (among all variants) and G671S (in the Delta variant) have an observed prevalence >15% among sequenced variant isolates (Supplemental Table 2). In contrast, multiple substitutions in the spike protein have been observed in all variants (Supplemental Figure 1 and Supplemental Table 3), which can result in immune evasion and reduced efficacy of monoclonal antibodies (43, 44). Structural analysis of the P323L, G671S, and F694Y (a highly prevalent substitution in early Omicron isolates) found each of these substitutions to be unlikely to reduce susceptibility to RDV. We confirmed RDV and GS-441524 retained antiviral activity against recombinant viruses containing each of these substitutions individually or in combinations. The findings were consistent with antiviral assessments performed in clinically isolated variants containing these Nsp12 substitutions, in which RDV and GS-441524 have similar potencies to WA1. Therefore, future variants containing P323L, G671S, F694Y, or their combinations are likely to remain susceptible to RDV and GS-441524.

Nsp12 mutations selected through in vitro passaging that are known to confer RDV resistance in coronaviruses (19, 35) are noticeably lacking from the sequence analysis of clinical samples. These mutations were observed in <0.001% of sequences analyzed, indicating that despite the widely prevalent use of RDV to treat COVID-19 in >10 million hospitalized patients over the course of the pandemic, emergence of RDV-resistant viruses is rare (42). However, with the recent expansion of RDV indication to treat COVID-19 earlier in the course of viral infection through outpatient use (11, 12) or potential future use of orally bioavailable prodrugs of GS- 441524, a sustained surveillance for emergence of resistance will need to continue.

In summary, we confirmed in several assay systems that past and present SARS-CoV-2 VOCs and VOIs retain *in vitro* susceptibility to both RDV and its parent nucleoside GS-441524. These findings highlight that both RDV and GS-441524 exhibit pan-variant SARS-CoV-2 activity and support the continued clinical use of RDV in approved patient populations.

## 4. Materials and Methods

### Reagents

Remdesivir (RDV) and GS-441524 were synthesized at Gilead Sciences, Inc. Validation of chemical identities were determined by NMR and LCMS, purity >95% was assessed by HPLC (7, 18). Compounds were solubilized in 100% dimethyl sulfoxide (DMSO) at a concentration of 10 mM.

### Viruses and cells

Vero-TMPRSS2 cells expressing human transmembrane serine protease 2 (hTMPRSS2) (45) were purchased from JCRB cell bank (Cat # JCRB 1818), National Institutes of Biomedical Innovation, Health and Nutrition. A549-ACE2 cells that stably express human angiotensin-converting enzyme 2 (hACE2) were established and provided by the University of Texas Medical Branch (46). A549-ACE2-TMPRSS2 (Cat # a549-hace2tpsa) were purchased from InvivoGen (San Diego, CA). All cells were maintained at 37°C and 5% CO_2_ in Dulbecco’s Minimum Essential Medium (DMEM) with GlutaMAX (Gibco cat # 10569-010) supplemented with 10% heat-inactivated fetal bovine serum (FBS) (Hyclone Cat # SH30396.03), 100 units/mL penicillin, 100 μg/mL streptomycin (Gibco Cat # 15140-122), and the appropriate selection agents - 1 mg/mL Geneticin (Vero-TMPRSS2), 10 μg/mL Blasticidin (A549-ACE2), or 0.5 μg/mL Puromycin and 100 μg/mL Hygromycin B (A549-ACE2-TMPRSS2). All cells were passaged 2-3 times per week with 0.25% Trypsin/0.02% EDTA (Gibco Cat#25200056). Cells used in all experimental set-ups were between passage 5 and 30.

SARS-CoV-2 isolates (Supplemental Table 1) were acquired through the World Reference Center for Emerging Viruses and Arboviruses at the University of Texas Medical Branch (Delta and Epsilon) and BEI Resources, National Institute of Allergy and Infectious Diseases (NIAID), National Institutes of Health (NIH). Isolates obtained from BEI Resources were deposited by the CDC (WA1 reference and Lambda), Bassam Hallis (Alpha), Alex Sigal and Tulio de Oliveira (Beta), the National Institute of Infectious Diseases (Gamma), Andrew S. Pekosz (Omicron and Zeta), Dr. Mehul Suthar and Dr. Benjamin Pinsky (Kappa), and Dr. David D. Ho (Iota).

All viruses were propagated 1-2 times in Vero-TMPRSS2 cells as follows. 1×10^7^ Vero- TMPRSS2 cells were seeded into a T225 flask in Vero-TMPRSS2 maintenance media and incubated overnight at 37°C + 5% CO_2_. The following day, the media was aspirated and replaced with 25 mL of DMEM supplemented with 2% FBS (infection medium) and infected with 10 μL of P0 stocks. The flasks were returned to 37°C + 5% CO_2_ until only 10-20% of viable cells remained (typically 36-72 hpi). The supernatant was harvested into a 50 mL Falcon tube and centrifuged at 2000 × *g* for 5 minutes to pellet cellular debris. The clarified supernatant was then transferred to a clean falcon tube and aliquoted as a working P1 stock into 100-250 μL aliquots and frozen at −80°C. The titer of the P1 stock was determined by plaque formation assay (PFA). If a second passage was required, the procedure above was repeated using the P1 stock to inoculate.

### Plaque formation assay (PFA)

3×10^5^ Vero-TMPRSS2 cells/well were seeded into 12-well plates in 1 mL of maintenance media and incubated overnight at 37°C and 5% CO_2_. The following day, cell confluency was confirmed to be >95% by visualization under a light microscope. Samples for analysis were serially diluted 10-fold in infection medium (DMEM + 2% FBS) up to a final dilution of 10^-5^ or 10^-6^. Spent supernatant was aspirated and replaced with 100 μL of serially diluted inoculum/well, and culture plates were returned to the incubator for 1 h with gentle rocking every 15 min. Following incubation, 2 mL of pre-warmed overlay medium (DMEM with 2% FBS, 1X penicillin/streptomycin, and 1.5% carboxymethylcellulose) was added to each well. Cells were then incubated without agitation for 3 days, at which point 2 mL of crystal violet fix/stain solution was added to each well. Cells were incubated at room temperature overnight.

Supernatants containing the crystal violet solution were discarded, and wells were washed with water 2 to 4 times each until plaques were visible and washes were clear of crystal violet residue. Plaques were counted manually from the most dilute wells consistently containing >5 plaque forming units (PFU).

### Plaque-reduction assay (PRA)

5×10^4^ A549-ACE2-TMPRSS2 cells were suspended into 500 μL maintenance medium and seeded into each well of a 48-well plate (Corning). Plates were incubated at 37°C with 5% CO_2_ overnight, after which the medium was aspirated and 250 μL of infection medium (DMEM + 2% FBS) was added to each well. Serial 3-fold dilutions of RDV in DMSO were added to each well using a Tecan D300e digital liquid dispenser. The DMSO concentrations were normalized to that of the highest compound concentration (DMSO less than <0.1% in final solution). SARS-CoV-2 was diluted into infection medium to 1×10^5^ PFU/mL, and 50 μL of inoculum was added to each well to result in a multiplicity of infection (MOI) = 0.1. At 48 or 72 hpi (for Omicron and 72 hpi WA1 reference), the supernatant was transferred to a clean 48-well plate and the plate sealed and frozen at −80°C until ready for analysis using the PFA assay described above. PFU counts for each variant were normalized to the DMSO controls for each variant (DMSO average = 0% inhibition). Due to the cumbersome nature of the PRA, all variants could not be read-out simultaneously; therefore, fold change calculations for this assay were assessed by taking the average EC_50_ for each variant divided by the average EC_50_ of the WA1 reference.

### Nucleoprotein ELISA

3×10^4^ A549-ACE2-TMPRSS2 cells in 100 μL DMEM (supplemented with 10% FBS and 1X penicillin/streptomycin) were seeded into each well of a 96-well plate and incubated overnight. The following day, media was aspirated and 100 μL of DMEM containing 2% FBS was added to each well. Three-fold serial dilutions of RDV or GS-441524 (in triplicate) were added to each well using a HP D300e digital dispenser with a final volume of 200 μL/well. Immediately after compound addition, cells were infected with 1.5×10^3^ PFU of the relevant SARS-CoV-2 variant diluted in 100 μL of DMEM supplemented with 2% FBS, resulting in a MOI = 0.05. Plates were centrifuged for 1 min at 500 × *g* and then incubated at 37°C with 5% CO_2_ for 2 days (or 3 days for Omicron strains and 72 hpi WA1 reference), after which media was aspirated, and cells fixed with 100% methanol for 10 minutes at room temperature (RT). The methanol was removed, and plates air-dried for 10 minutes at RT followed by a 1 h incubation with 100 μL/well of blocking buffer (phosphate-buffered saline [PBS] with 10% FBS, 5% non-fat dry milk, and 0.1% Tween 20) for 1 h at 37°C. The blocking buffer was then aspirated and 50 μL of a 1:4000 dilution of rabbit anti-SARS-CoV-2 nucleocapsid (N) antibody (MA536086, Invitrogen) in blocking buffer was added and incubated for 2 h at 37°C. Plates were washed 4× with 200 μL/well of PBS containing 0.1% Tween 20 prior to addition of 50 μL/well of horseradish peroxidase (HRP) conjugated goat-anti-rabbit IgG (GtxRb-003-FHRPX, ImmunoReagents) diluted 1:4000 in blocking buffer. Plates were again incubated for 1 h at 37°C and then washed 4× with 200 μL PBS with 0.1% Tween 20. 100 μL TMB reagent (ENN301, Thermo Scientific) was added to each well and allowed to incubate at RT until visible staining of the positive control wells, usually 5-10 minutes. The reaction was stopped with addition of 100 μL/well of TMB stop solution (5150-0021 SeraCare). The absorbance was then read at 450 nm using an EnVision plate reader. Fold change for variants was calculated for each experiment, comparing to the relevant WA1 reference. Fold change across all experiments was then averaged to obtain final reported values.

### SARS-CoV-2 sequence analysis

The tabulated amino acid substitutions from WA1 reference (MN985325) for a total of 5,842,948 SARS-Cov-2 genome sequences were obtained from GISAID EpiCov database as of January 18, 2021 (https://www.gisaid.org/) (47). Sequences with length <29,000 nucleotides in length or that contained >5% of ambiguous bases across genome were excluded from analyses. The sequences were further categorized into 11 VOC/VOIs according to the PANGO lineage using Pangolin software (48). The Regeneron COVID-19 Dashboard web portal (https://covid19dashboard.regeneron.com) was used to assess the overall prevalence of mutations in 7,106,062 unfiltered sequences from GISAID database on January 18, 2022. Lineage-associated amino acid changes were obtained from PANGO lineage web portal (https://cov-lineages.org/).

### Protein structure modelling and visualization

The model of pre-incorporated RDV-TP in the active site of the SARS-CoV-2 polymerase complex was developed from the NTP-free cryo-EM structure 6XEZ and has been described elsewhere (37). The variant mutations P323L, P323L/G671S, and P323L/F694Y were introduced and optimized by conducting a side chain rotamer optimization and minimization of the mutated residues and surrounding residues within 5 Å using Prime. The impact of each mutation on the predicted binding affinity to RDV-TP was assessed with an MM-GBSA residue scan within Bioluminate.

### Site-directed mutagenesis and recombinant virus rescue

To produce recombinant SARS-CoV-2 virus, we utilized a SARS-CoV-2 reverse genetics system previously described (24, 49) that was slightly modified by fusing plasmids F1-F3 single plasmid making it a 3-plasmid reverse genetics system producing infectious virus containing either Nano luciferase (Nluc) or the Firefly luciferase (Fluc) transgene. Desired substitutions in nsp12 of the SARS-CoV-2 genome were added to the nsp12 containing F4 plasmid using the Quick-Change PCR protocol using Platinum SuperFI II PCR master-mix (ThermoFisher Scientific cat. No. 12361010) following manufacturer’s protocols. The primers used to engineer specific mutations were SARS_CoV2_NSP12_P323L_Fw-5’-GTTCCCACTTACAAGTTTTG-3’ and SARS_CoV2_NSP12_P323L_Rv- 5’-CAAAACTTGTAAGTGGGAAC-3’ for P323L, SARS_CoV2_NSP12_F694Y_Fw-5’-GCTAATAGTGTTTATAACATTTGTC-3’ and SARS_CoV2_NSP12_F694Y_Rv-5’-GACAAATGTTATAAACACTATTAGC-3’ for F694Y, and SARS_CoV2_NSP12_G671S_Fw-5’-GTCATGTGTGGCAGTTCACTATATG-3’ and SARS_CoV2_NSP 12_G671S_Rv-5 ‘-CATATAGTGAACTGCCACACATGAC-3’ for G671S. Substitutions (red highlights in primers) were sequenced confirmed, and then validated plasmids were digested with either BsaI or Esp3I. Cut plasmids were then ligated together using T4 DNA ligase, and the ligated product was *in vitro* transcribed into RNA. The RNA products were then electroporated into Vero-TMPRSS2 cells and monitored until extensive cytopathic effect was observed and P0 virus harvested. P0 virus stocks were titered and passaged to P1 as described above for propagation of clinical isolates. Virus used for experiments was either P1 (Fluc) or P2 (Nluc).

### Construction of a recombinant Omicron SARS-CoV-2

Recombinant Omicron SARS-CoV-2 was constructed by engineering the complete mutations from Omicron variant (GISAID EPI_ISL_6640916) into an infectious cDNA clone of clinical isolate USA-WA1/2020 (50). All mutations were introduced into the infectious cDNA clone of USA-WA1/2020 using PCR-based mutagenesis as previously described (51). An additional recombinant Omicron SARS-CoV-2 was generated bearing the F694Y substitution in NSP12 by the methods detailed above. rOmicron viruses were analyzed using the N-protein ELISA following the protocol used for clinical isolates.

### Antiviral activity assessment from recombinant luciferase containing viruses

For Nluc readouts, 1.2×10^4^ A549-hACE2 cells per well were suspended in 50 μL infection medium and seeded into a white clear-bottom 96-well plate (Corning) and incubated overnight at 37°C with 5% CO_2_. On the following day, compounds were added directly to cultures as 3-fold serial dilutions with a Tecan D300e digital liquid dispenser, with DMSO volumes normalized to that of the highest compound concentration (final DMSO concentration <0.1%). SARS-CoV-2- Nluc viruses were diluted to MOI = 0.05 and aliquoted 50 μL/well. At 48 hpi, 75 μL Nluc substrate solution (Promega) was added to each well. Luciferase signals were measured using an Envision microplate reader (Perkin Elmer).

For Firefly luciferase readouts, the assay set-up was the same as the Nluc assay except cells were infected with SARS-CoV-2-Fluc viruses at an MOI = 1.0 and at 48 hpi, 100 μL One-Glo luciferase substrate solution (Promega) was added to each well prior to reading the signal on the Envision plate reader (Perkin Elmer).

### EC_50_ determinations

The half-maximal effective concentration (EC_50_) is defined as the compound concentration at which there was a 50% reduction in plaque formation (PRA), luciferase signal, or N-protein expression (ELISA) relative to infected cells with DMSO alone (0% inhibition) and uninfected control (100% inhibition). EC_50_ values were determined using GraphPad Prism 8.1.2 using non-linear regression curve fits. Constraints were used when required to ensure the bottom or top of the fit curves were close to 0 and 100, respectively.

## 5. Acknowledgments

Becky Norquist provided comprehensive manuscript writing support. We thank Kenneth S. Plante, Jessica A. Plante, and David S. Blakeman for coordination of virus stocks from the World Reference Center for Emerging Viruses and Arboviruses at the University of Texas Medical Branch. These studies were fully funded by Gilead Sciences, Inc.

## Supplemental Materials

**Supplemental Figure 1.**
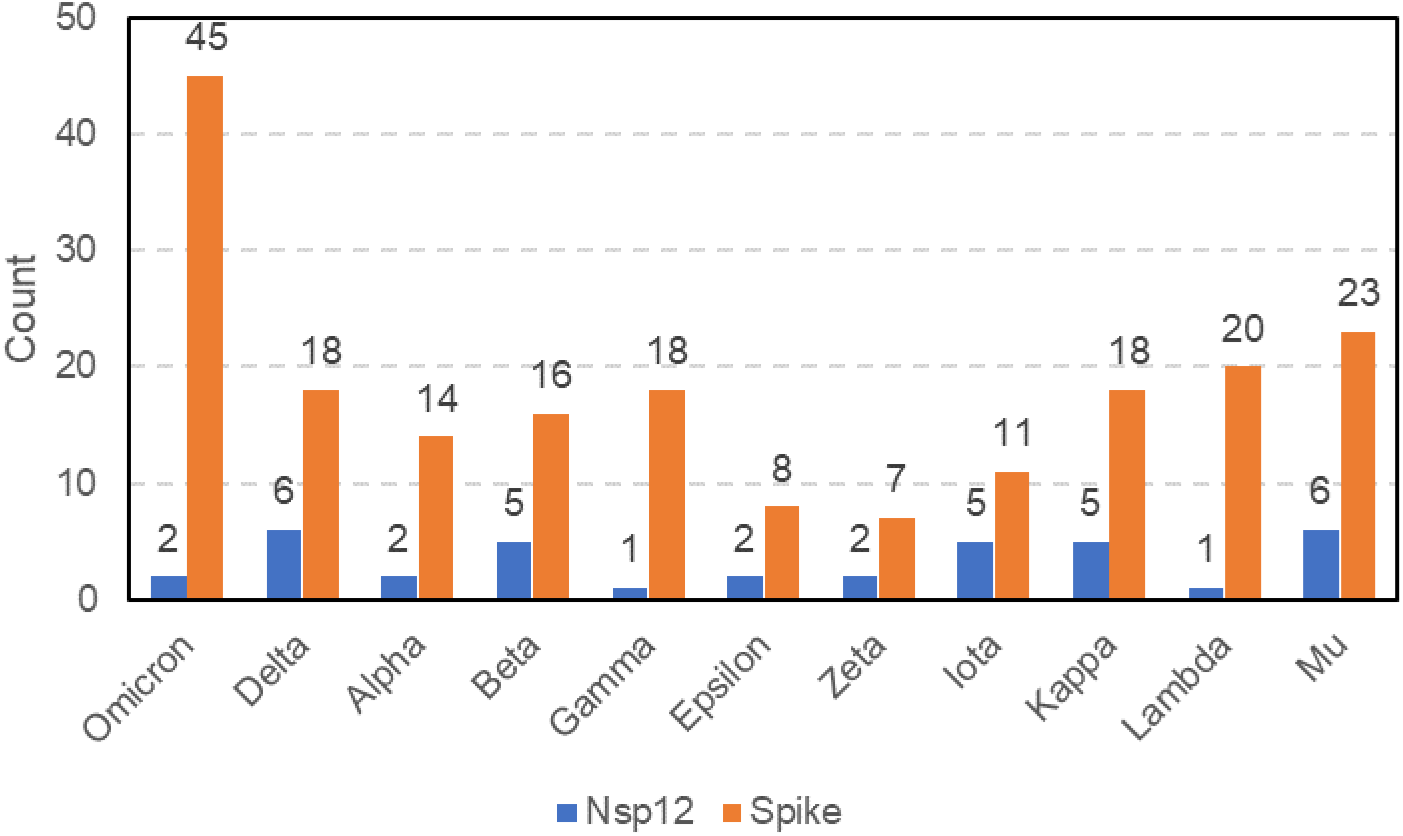
Amino acid substitutions in Nsp12 and Spike. The total number of amino acid substitutions in Nsp12 compared to Spike from each variant of concern or variant of interest. Actual substitutions for each variant are found in Supplemental Tables 1 (Nsp12) and 2 (Spike).

**Supplemental Figure 2.**
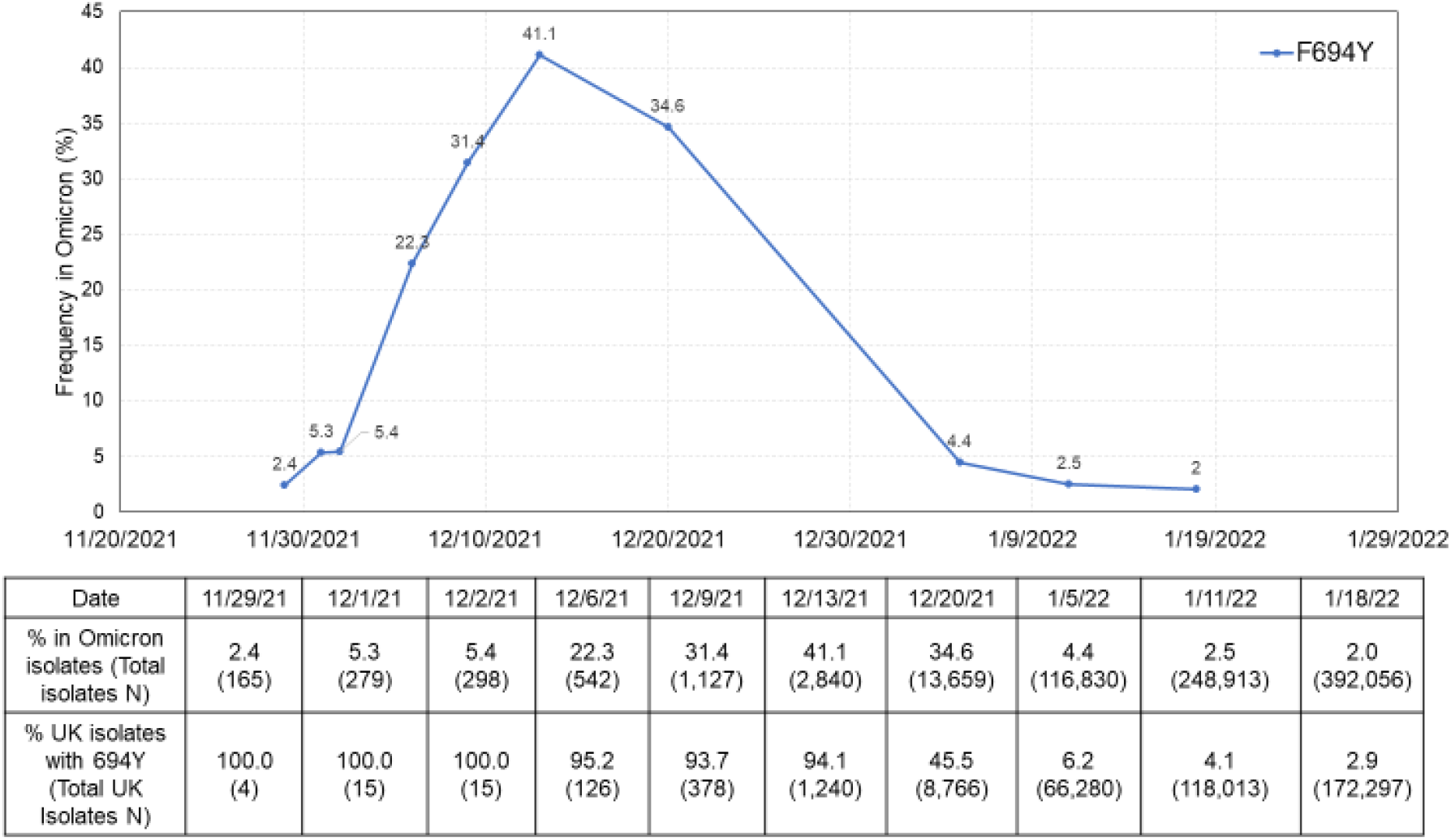
F694Y frequency in Omicron variant over time. Omicron sequences were obtained from GISAID at ten different timepoints. Frequency of F694Y in all omicron sequences at each timepoint are plotted. The frequency among UK isolates is shown in the data table below the plot.

**Supplemental Figure 3.**
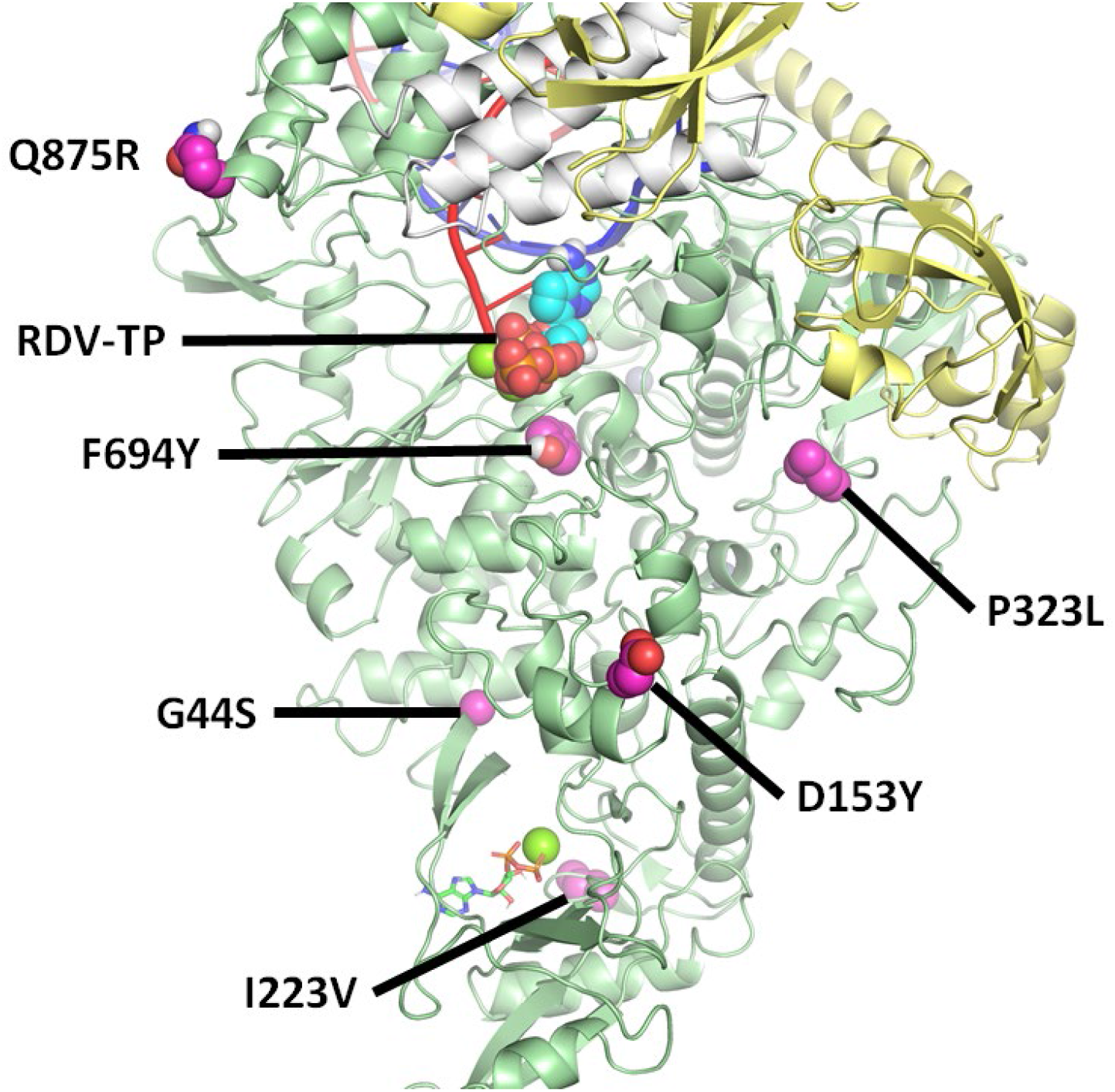
Location of Omicron Nsp12 amino acid substitutions with frequency >0.5%. Amino acid substitutions identified in Table 2 are mapped onto the model of the polymerase complex with pre-incorporated RDV-TP. All substitutions occur on the surface of Nsp12, away from the active site, apart from F694Y. Nsp12 is shown in green, Nsp8 in yellow, Nsp7 in white, template RNA in blue, and primer RNA in red.

**Supplemental Figure 4.**
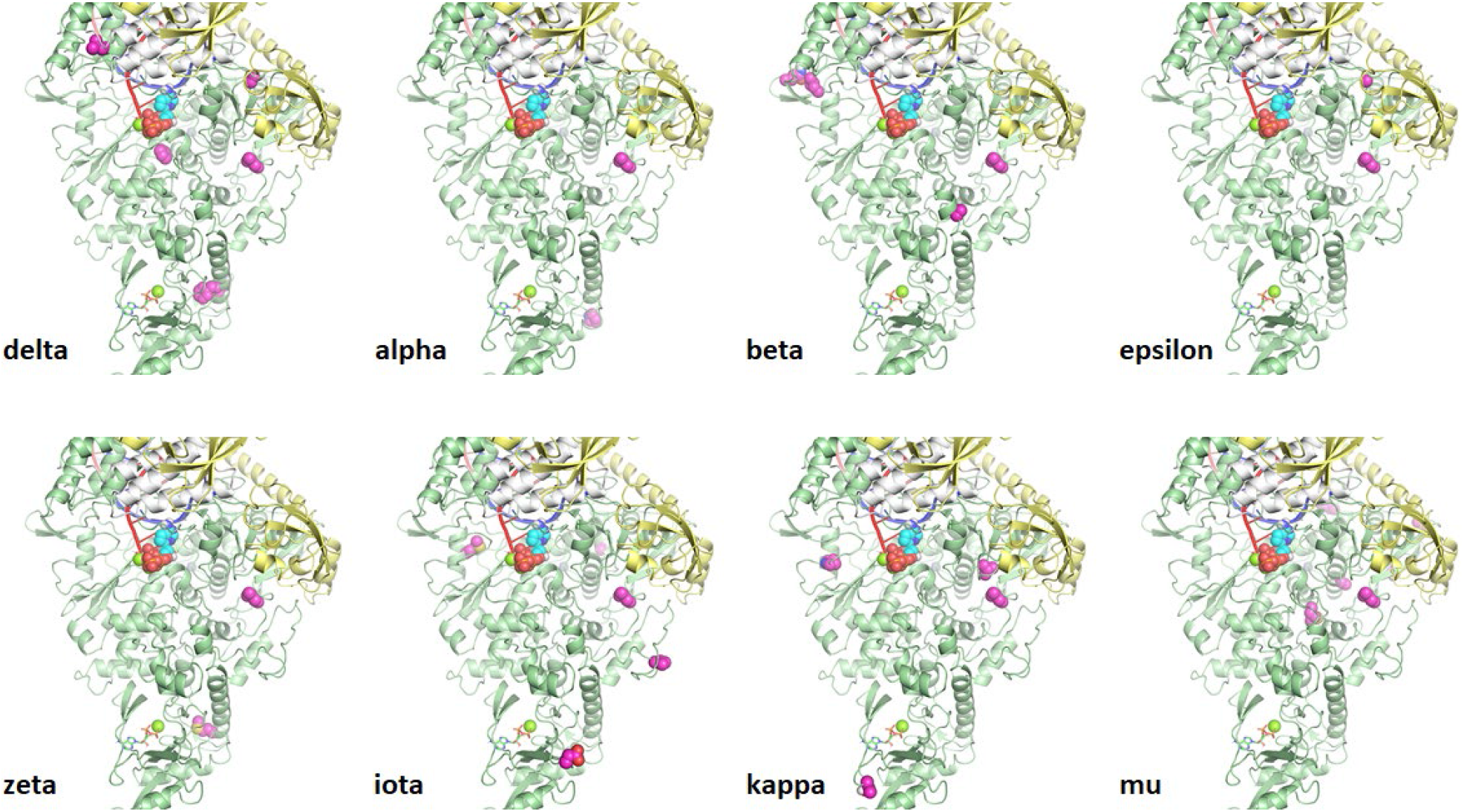
Location of variant Nsp12 amino acid substitutions with frequency >1%. Amino acid substitutions identified in Supplemental Table 1 are mapped onto the model of the polymerase complex with pre-incorporated RDV-TP. Gamma and Lambda are not shown, since substitutions are limited to P323L, while Omicron substitutions are shown in Supplemental Fig. 3. With the exception of F694Y, seen in Delta, the substitutions are all remote from the polymerase active site. Nsp12 is shown in green, Nsp8 in yellow, Nsp7 in white, template RNA in blue, and primer RNA in red. Amino acid substitutions are shown in magenta.

**Supplemental Table 1.**
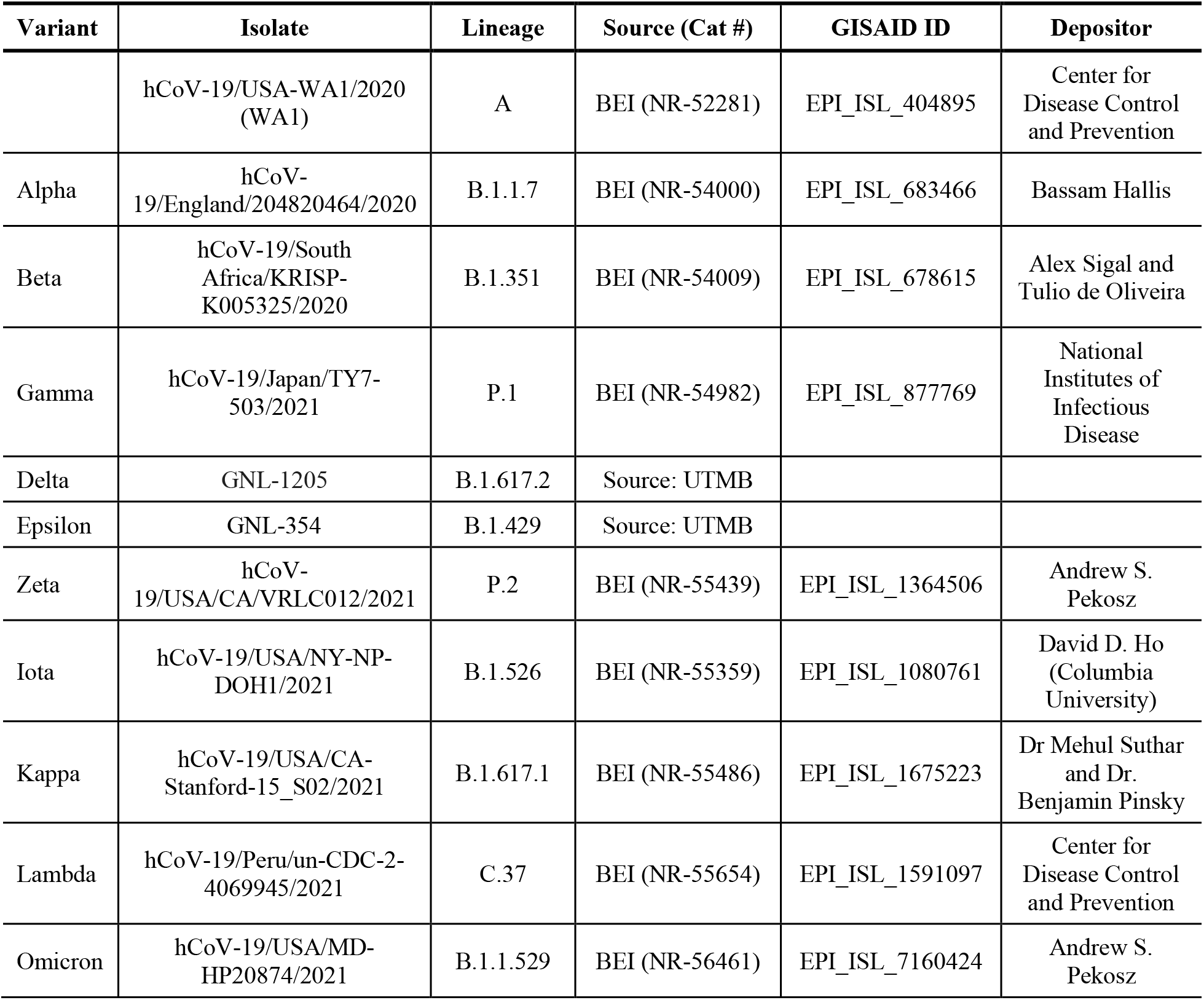
SARS-CoV-2 variant clinical isolates and sources.

**Supplemental Table 2.** Amino acid substitutions of Nsp12 in VOC/VOI observed at frequency ≥1%.

| **SARS-CoV-2 Variant** | **SARS-CoV-2 Variant Sequences Analyzed (N, % of Total Sequences)** | **Nsp12 Substitution** | **Frequency of Substitution, % (N)** |
| --- | --- | --- | --- |
| Omicron | 392,056 (6.7%) | P323L | 99.5 (390,020) |
|  |  | F694Y | 2.0 (7,822) |
| Delta | 4,059,836 (69.5%) | P323L | 99.4 (4,034,234) |
|  |  | G671S | 97.8 (3,968,940) |
|  |  | L838I | 6.8 (277,419) |
|  |  | F192V | 5.8 (235,631) |
|  |  | F694Y | 4.9 (197,489) |
|  |  | R197Q | 1.2 (47,122) |
| Alpha | 1,158,351 (19.8%) | P323L | 99.3 (1,149,857) |
|  |  | P227L | 14.2 (164,221) |
| Beta | 35,180 (0.60%) | P323L | 91.7 (37,661) |
|  |  | L829F | 9.0 (3,712) |
|  |  | P323H | 7.5 (3,075) |
|  |  | Q822H | 5.4 (2,198) |
|  |  | A176T | 2.3 (940) |
| Gamma | 120,614 (2.1%) | P323L | 99.3 (119,734) |
| Epsilon | 44,549 (0.76%) | P323L | 94.9 (42,274) |
|  |  | G671V | 2.1 (937) |
| Zeta | 1,834 (0.03%) | P323L | 98.2 (1,801) |
|  |  | M196L | 1.9 (34) |
| Iota | 20,923 (0.36%) | P323L | 99.5 (20,821) |
|  |  | M601I | 2.3 (485) |
|  |  | V257A | 2.1 (438) |
|  |  | D62G | 1.2 (244) |
|  |  | A529V | 1.2 (241) |
| Kappa | 5,498 (0.09%) | P323L | 99.5 (5,472) |
|  |  | V675I | 7.8 (431) |
|  |  | K478N | 7.1 (388) |
|  |  | T26I | 4.0(218) |
|  |  | P809R | 1.6 (88) |
| Lambda | 6,219 (0.11%) | P323L | 99.8 (6,209) |
| Mu | 7,888 (0.14%) | P323L | 98.5 (7,771) |
|  |  | Y521C | 21.8 (1,722) |
|  |  | M629I | 9.6 (756) |
|  |  | T344I | 2.3 (181) |
|  |  | A656S | 1.6 (124) |
|  |  | S364F | 1.4 (109) |

**Supplemental Table 3.**
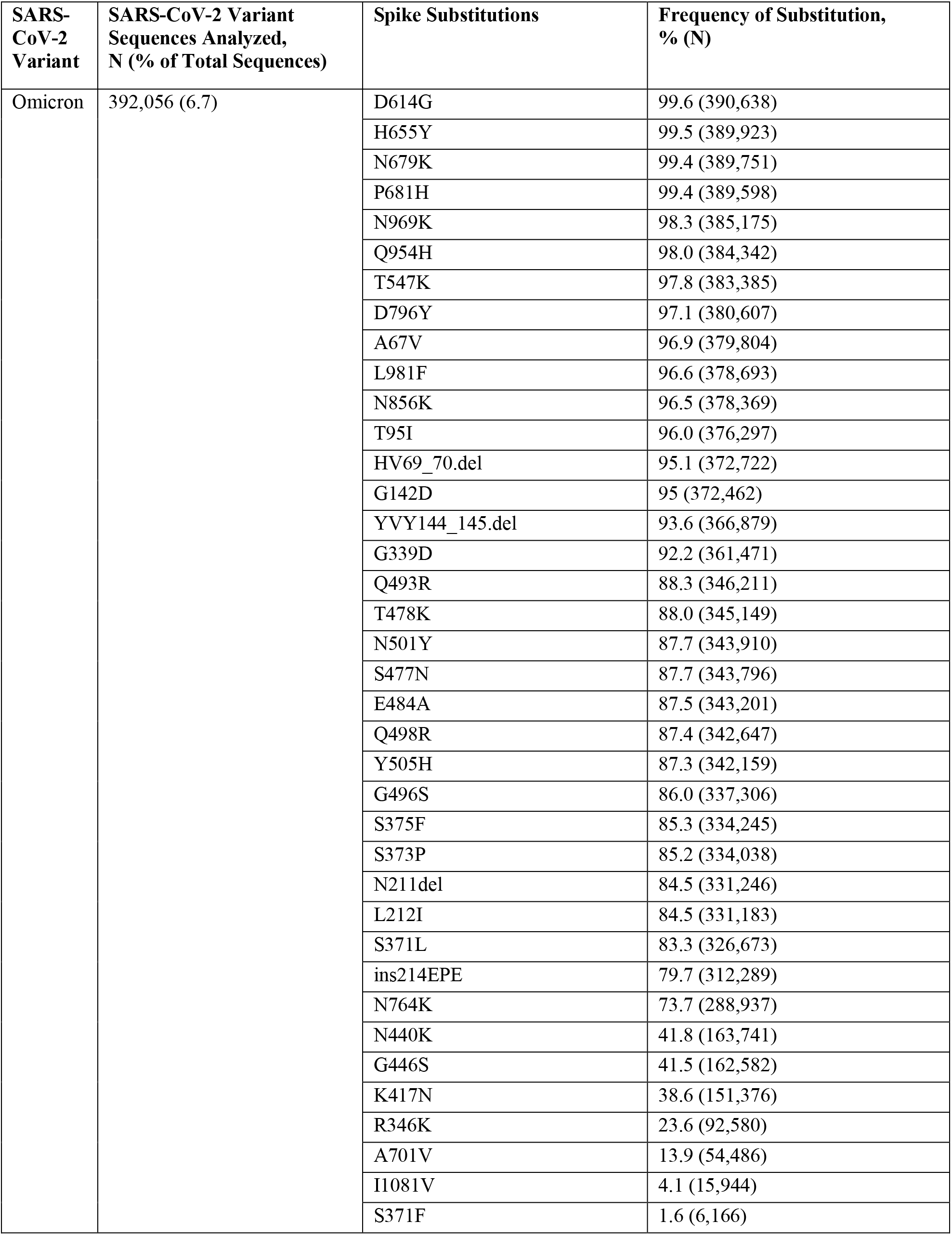
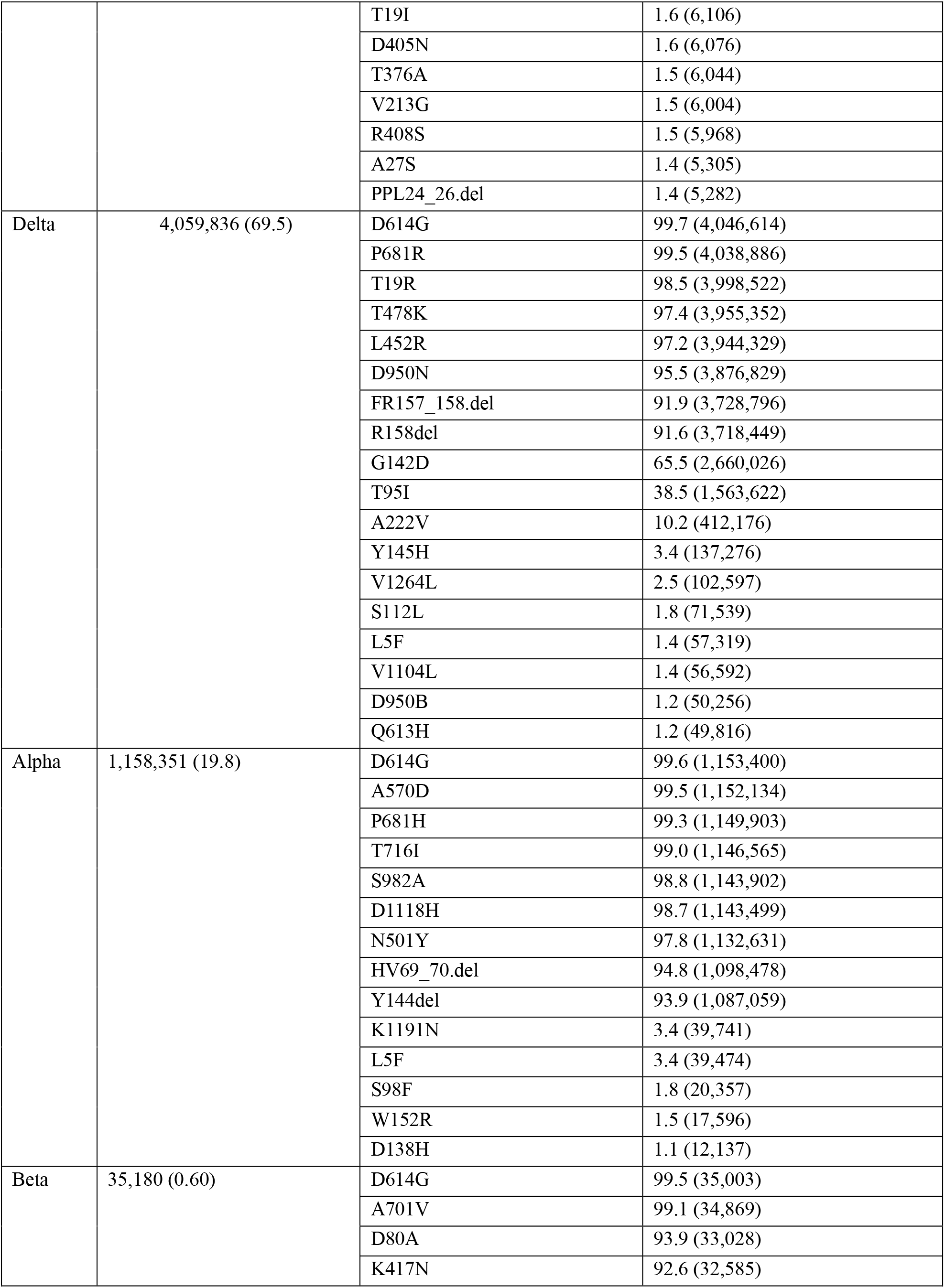
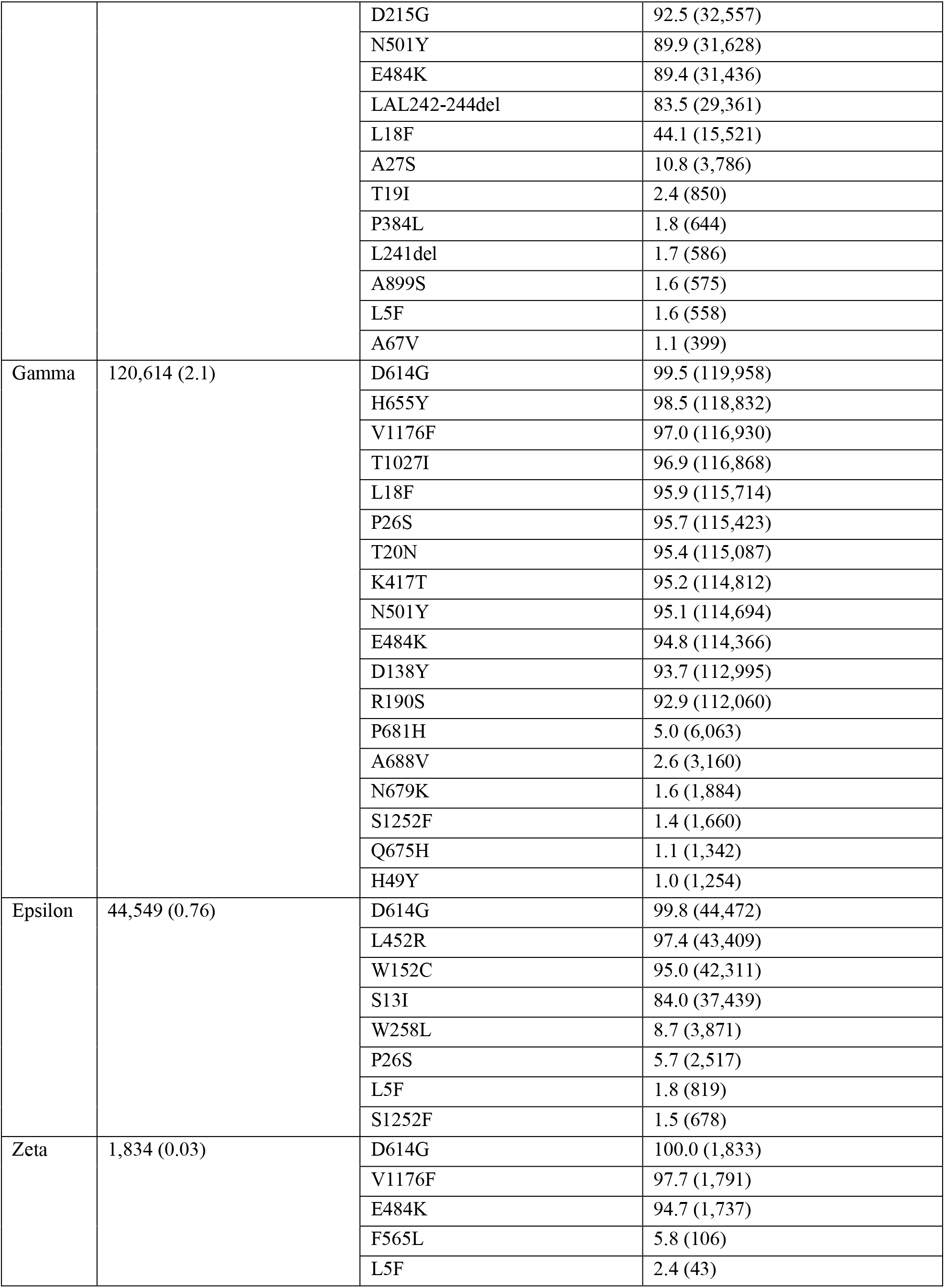
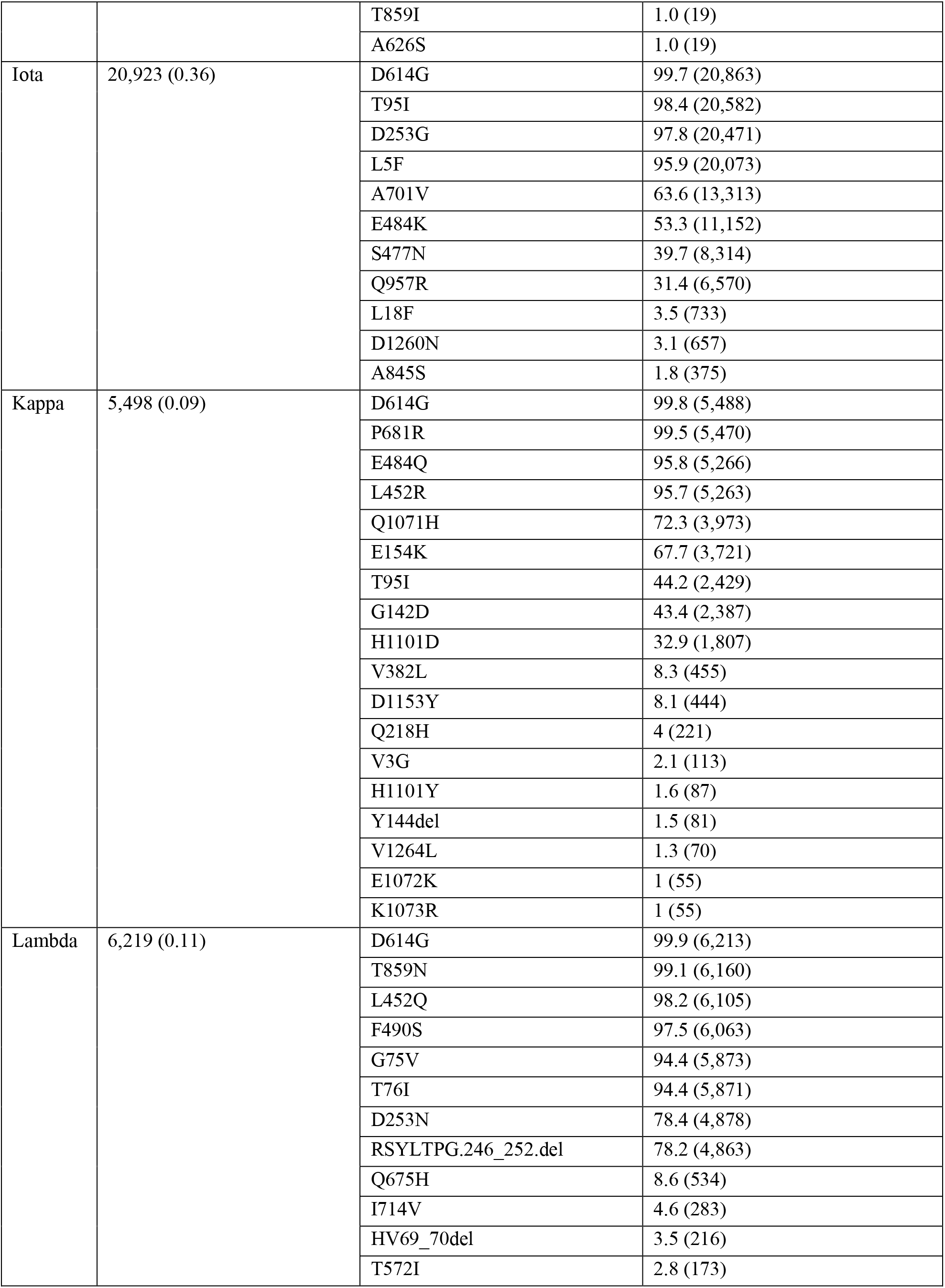
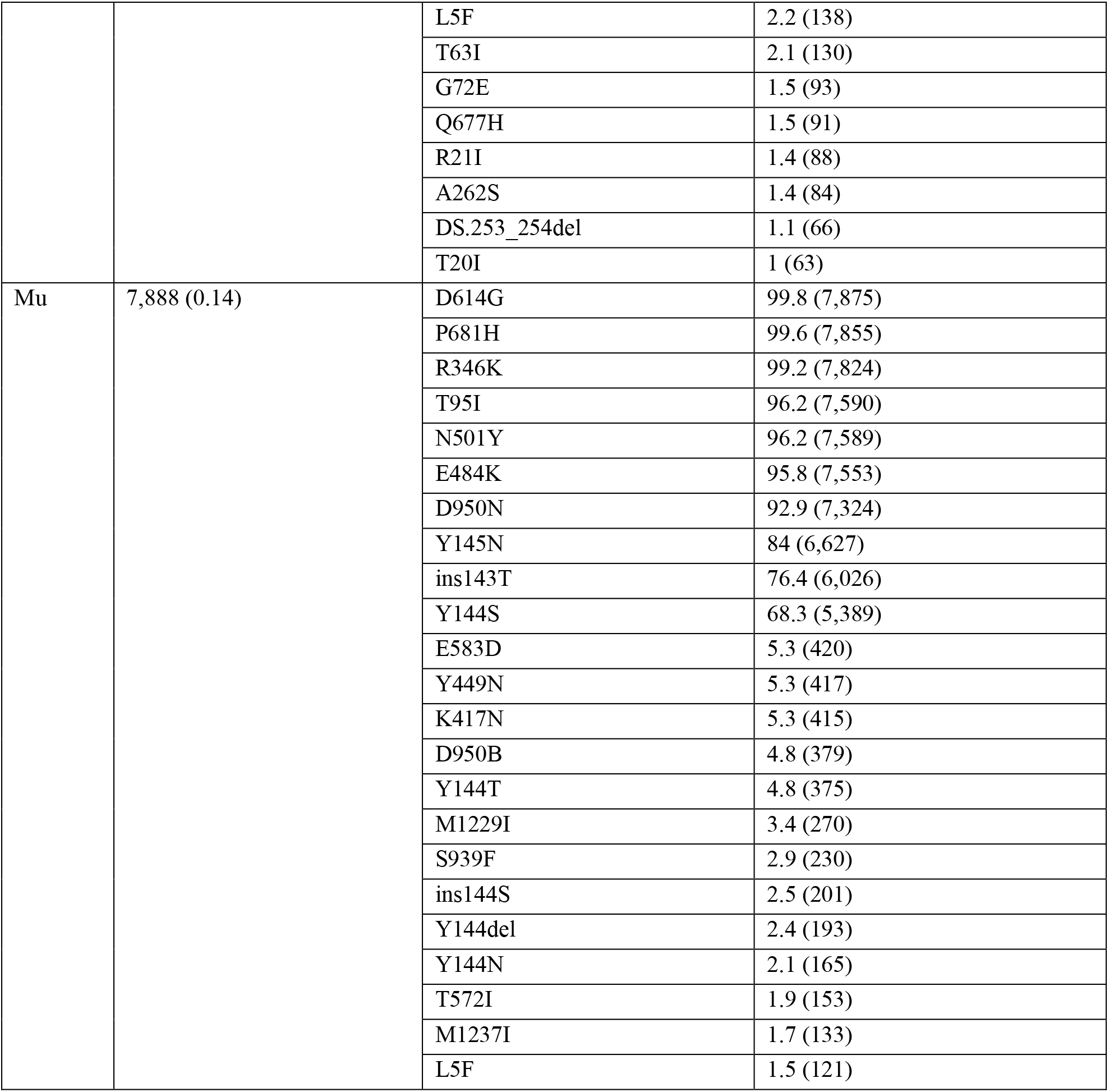
Amino acid substitutions of Spike in VOC/VOI observed at frequency ≥1%.

| SARS-CoV-2 Variant | SARS-CoV-2 Variant Sequences Analyzed, N (% of Total Sequences) | Spike Substitutions | Frequency of Substitution, % (N) |
| --- | --- | --- | --- |
| Omicron | 392,056 (6.7) | D614G | 99.6 (390,638) |
|  |  | H655Y | 99.5 (389,923) |
|  |  | N679K | 99.4 (389,751) |
|  |  | P681H | 99.4 (389,598) |
|  |  | N969K | 98.3 (385,175) |
|  |  | Q954H | 98.0 (384,342) |
|  |  | T547K | 97.8 (383,385) |
|  |  | D796Y | 97.1 (380,607) |
|  |  | A67V | 96.9 (379,804) |
|  |  | L981F | 96.6 (378,693) |
|  |  | N856K | 96.5 (378,369) |
|  |  | T95I | 96.0 (376,297) |
|  |  | HV69_70.del | 95.1 (372,722) |
|  |  | G142D | 95 (372,462) |
|  |  | YVY144_145.del | 93.6 (366,879) |
|  |  | G339D | 92.2 (361,471) |
|  |  | Q493R | 88.3 (346,211) |
|  |  | T478K | 88.0 (345,149) |
|  |  | N501Y | 87.7 (343,910) |
|  |  | S477N | 87.7 (343,796) |
|  |  | E484A | 87.5 (343,201) |
|  |  | Q498R | 87.4 (342,647) |
|  |  | Y505H | 87.3 (342,159) |
|  |  | G496S | 86.0 (337,306) |
|  |  | S375F | 85.3 (334,245) |
|  |  | S373P | 85.2 (334,038) |
|  |  | N211del | 84.5 (331,246) |
|  |  | L212I | 84.5 (331,183) |
|  |  | S371L | 83.3 (326,673) |
|  |  | ins214EPE | 79.7 (312,289) |
|  |  | N764K | 73.7 (288,937) |
|  |  | N440K | 41.8 (163,741) |
|  |  | G446S | 41.5 (162,582) |
|  |  | K417N | 38.6 (151,376) |
|  |  | R346K | 23.6 (92,580) |
|  |  | A701V | 13.9 (54,486) |
|  |  | I1081V | 4.1 (15,944) |
|  |  | S371F | 1.6 (6,166) |
|  |  | T19I | 1.6 (6,106) |
|  |  | D405N | 1.6 (6,076) |
|  |  | T376A | 1.5 (6,044) |
|  |  | V213G | 1.5 (6,004) |
|  |  | R408S | 1.5 (5,968) |
|  |  | A27S | 1.4 (5,305) |
|  |  | PPL24_26.del | 1.4 (5,282) |
| Delta | 4,059,836 (69.5) | D614G | 99.7 (4,046,614) |
|  |  | P681R | 99.5 (4,038,886) |
|  |  | T19R | 98.5 (3,998,522) |
|  |  | T478K | 97.4 (3,955,352) |
|  |  | L452R | 97.2 (3,944,329) |
|  |  | D950N | 95.5 (3,876,829) |
|  |  | FR157_158.del | 91.9 (3,728,796) |
|  |  | R158del | 91.6 (3,718,449) |
|  |  | G142D | 65.5 (2,660,026) |
|  |  | T95I | 38.5 (1,563,622) |
|  |  | A222V | 10.2 (412,176) |
|  |  | Y145H | 3.4 (137,276) |
|  |  | V1264L | 2.5 (102,597) |
|  |  | S112L | 1.8 (71,539) |
|  |  | L5F | 1.4 (57,319) |
|  |  | V1104L | 1.4 (56,592) |
|  |  | D950B | 1.2 (50,256) |
|  |  | Q613H | 1.2 (49,816) |
| Alpha | 1,158,351 (19.8) | D614G | 99.6 (1,153,400) |
|  |  | A570D | 99.5 (1,152,134) |
|  |  | P681H | 99.3 (1,149,903) |
|  |  | T716I | 99.0 (1,146,565) |
|  |  | S982A | 98.8 (1,143,902) |
|  |  | D1118H | 98.7 (1,143,499) |
|  |  | N501Y | 97.8 (1,132,631) |
|  |  | HV69_70.del | 94.8 (1,098,478) |
|  |  | Y144del | 93.9 (1,087,059) |
|  |  | K1191N | 3.4 (39,741) |
|  |  | L5F | 3.4 (39,474) |
|  |  | S98F | 1.8 (20,357) |
|  |  | W152R | 1.5 (17,596) |
|  |  | D138H | 1.1 (12,137) |
| Beta | 35,180 (0.60) | D614G | 99.5 (35,003) |
|  |  | A701V | 99.1 (34,869) |
|  |  | D80A | 93.9 (33,028) |
|  |  | K417N | 92.6 (32,585) |
|  |  | D215G | 92.5 (32,557) |
|  |  | N501Y | 89.9 (31,628) |
|  |  | E484K | 89.4 (31,436) |
|  |  | LAL242-244del | 83.5 (29,361) |
|  |  | L18F | 44.1 (15,521) |
|  |  | A27S | 10.8 (3,786) |
|  |  | T19I | 2.4 (850) |
|  |  | P384L | 1.8 (644) |
|  |  | L241del | 1.7 (586) |
|  |  | A899S | 1.6 (575) |
|  |  | L5F | 1.6 (558) |
|  |  | A67V | 1.1 (399) |
| Gamma | 120,614 (2.1) | D614G | 99.5 (119,958) |
|  |  | H655Y | 98.5 (118,832) |
|  |  | V1176F | 97.0 (116,930) |
|  |  | T1027I | 96.9 (116,868) |
|  |  | L18F | 95.9 (115,714) |
|  |  | P26S | 95.7 (115,423) |
|  |  | T20N | 95.4 (115,087) |
|  |  | K417T | 95.2 (114,812) |
|  |  | N501Y | 95.1 (114,694) |
|  |  | E484K | 94.8 (114,366) |
|  |  | D138Y | 93.7 (112,995) |
|  |  | R190S | 92.9 (112,060) |
|  |  | P681H | 5.0 (6,063) |
|  |  | A688V | 2.6 (3,160) |
|  |  | N679K | 1.6 (1,884) |
|  |  | S1252F | 1.4 (1,660) |
|  |  | Q675H | 1.1 (1,342) |
|  |  | H49Y | 1.0 (1,254) |
| Epsilon | 44,549 (0.76) | D614G | 99.8 (44,472) |
|  |  | L452R | 97.4 (43,409) |
|  |  | W152C | 95.0 (42,311) |
|  |  | S13I | 84.0 (37,439) |
|  |  | W258L | 8.7 (3,871) |
|  |  | P26S | 5.7 (2,517) |
|  |  | L5F | 1.8 (819) |
|  |  | S1252F | 1.5 (678) |
| Zeta | 1,834 (0.03) | D614G | 100.0 (1,833) |
|  |  | V1176F | 97.7 (1,791) |
|  |  | E484K | 94.7 (1,737) |
|  |  | F565L | 5.8 (106) |
|  |  | L5F | 2.4 (43) |
|  |  | T859I | 1.0 (19) |
|  |  | A626S | 1.0 (19) |
| Iota | 20,923 (0.36) | D614G | 99.7 (20,863) |
|  |  | T95I | 98.4 (20,582) |
|  |  | D253G | 97.8 (20,471) |
|  |  | L5F | 95.9 (20,073) |
|  |  | A701V | 63.6 (13,313) |
|  |  | E484K | 53.3 (11,152) |
|  |  | S477N | 39.7 (8,314) |
|  |  | Q957R | 31.4 (6,570) |
|  |  | L18F | 3.5 (733) |
|  |  | D1260N | 3.1 (657) |
|  |  | A845S | 1.8 (375) |
| Kappa | 5,498 (0.09) | D614G | 99.8 (5,488) |
|  |  | P681R | 99.5 (5,470) |
|  |  | E484Q | 95.8 (5,266) |
|  |  | L452R | 95.7 (5,263) |
|  |  | Q1071H | 72.3 (3,973) |
|  |  | E154K | 67.7 (3,721) |
|  |  | T95I | 44.2 (2,429) |
|  |  | G142D | 43.4 (2,387) |
|  |  | H1101D | 32.9 (1,807) |
|  |  | V382L | 8.3 (455) |
|  |  | D1153Y | 8.1 (444) |
|  |  | Q218H | 4 (221) |
|  |  | V3G | 2.1 (113) |
|  |  | H1101Y | 1.6 (87) |
|  |  | Y144del | 1.5 (81) |
|  |  | V1264L | 1.3 (70) |
|  |  | E1072K | 1 (55) |
|  |  | K1073R | 1 (55) |
| Lambda | 6,219 (0.11) | D614G | 99.9 (6,213) |
|  |  | T859N | 99.1 (6,160) |
|  |  | L452Q | 98.2 (6,105) |
|  |  | F490S | 97.5 (6,063) |
|  |  | G75V | 94.4 (5,873) |
|  |  | T76I | 94.4 (5,871) |
|  |  | D253N | 78.4 (4,878) |
|  |  | RSYLTPG.246_252.del | 78.2 (4,863) |
|  |  | Q675H | 8.6 (534) |
|  |  | I714V | 4.6 (283) |
|  |  | HV69_70del | 3.5 (216) |
|  |  | T572I | 2.8 (173) |
|  |  | L5F | 2.2 (138) |
|  |  | T63I | 2.1 (130) |
|  |  | G72E | 1.5 (93) |
|  |  | Q677H | 1.5 (91) |
|  |  | R21I | 1.4 (88) |
|  |  | A262S | 1.4 (84) |
|  |  | DS.253_254del | 1.1 (66) |
|  |  | T20I | 1 (63) |
| Mu | 7,888 (0.14) | D614G | 99.8 (7,875) |
|  |  | P681H | 99.6 (7,855) |
|  |  | R346K | 99.2 (7,824) |
|  |  | T95I | 96.2 (7,590) |
|  |  | N501Y | 96.2 (7,589) |
|  |  | E484K | 95.8 (7,553) |
|  |  | D950N | 92.9 (7,324) |
|  |  | Y145N | 84 (6,627) |
|  |  | ins143T | 76.4 (6,026) |
|  |  | Y144S | 68.3 (5,389) |
|  |  | E583D | 5.3 (420) |
|  |  | Y449N | 5.3 (417) |
|  |  | K417N | 5.3 (415) |
|  |  | D950B | 4.8 (379) |
|  |  | Y144T | 4.8 (375) |
|  |  | M1229I | 3.4 (270) |
|  |  | S939F | 2.9 (230) |
|  |  | ins144S | 2.5 (201) |
|  |  | Y144del | 2.4 (193) |
|  |  | Y144N | 2.1 (165) |
|  |  | T572I | 1.9 (153) |
|  |  | M1237I | 1.7 (133) |
|  |  | L5F | 1.5 (121) |

## References

1. Eckerle LD, Becker MM, Halpin RA, Li K, Venter E, Lu X, Scherbakova S, Graham RL, Baric RS, Stockwell TB, Spiro DJ, Denison MR. 2010. Infidelity of SARS-CoV Nsp14-exonuclease mutant virus replication is revealed by complete genome sequencing. PLoS Pathog 6:e1000896.

2. Gribble J, Stevens LJ, Agostini ML, Anderson-Daniels J, Chappell JD, Lu X, Pruijssers AJ, Routh AL, Denison MR. 2021. The coronavirus proofreading exoribonuclease mediates extensive viral recombination. PLoS Pathog 17:e1009226

3. Washington NL, Gangavarapu K, Zeller M, Bolze A, Cirulli ET, Schiabor Barrett KM, Larsen BB, Anderson C, White S, Cassens T, Jacobs S, Levan G, Nguyen J, Ramirez JM 3rd, Rivera-Garcia C, Sandoval E, Wang X, Wong D, Spencer E, Robles-Sikisaka R, Kurzban E, Hughes LD, Deng X, Wang C, Servellita V, Valentine H, De Hoff P, Seaver P, Sathe S, Gietzen K, Sickler B, Antico J, Hoon K, Liu J, Harding A, Bakhtar O, Basler T, Austin B, MacCannell D, Isaksson M, Febbo PG, Becker D, Laurent M, McDonald E, Yeo GW, Knight R, Laurent LC, de Feo E, Worobey M, Chiu CY, Suchard MA, Lu JT, Lee W, Andersen KG. 2021. Emergence and rapid transmission of SARS-CoV-2 B.1.1.7 in the United States. Cell 184:2587–2594.e7.

4. Mishra S, Mindermann S, Sharma M, Whittaker C, Mellan TA, Wilton T, Klapsa D, Mate R, Fritzsche M, Zambon M, Ahuja J, Howes A, Miscouridou X, Nason GP, Ratmann O, Semenova E, Leech G, Sandkühler JF, Rogers-Smith C, Vollmer M, Unwin HJT, Gal Y, Chand M, Gandy A, Martin J, Volz E, Ferguson NM, Bhatt S, Brauner JM, Flaxman S; COVID-19 Genomics UK (COG-UK) Consortium. 2021. Changing composition of SARS-CoV-2 lineages and rise of Delta variant in England. EClinicalMedicine 39:101064.

5. Sah P, Vilches TN, Shoukat A, Fitzpatrick MC, Pandey A, Singer BH, Moghadas SM, Galvani AP. 2021. Quantifying the potential dominance of immune-evading SARS-CoV-2 variants in the United States. medRxiv 2021.05.10.21256996.

6. European Centre for Disease Prevention and Control (ECDC). 2021. Threat Assessment Brief: Implications of the emergence and spread of the SARS-CoV-2 B.1.1. 529 variant of concern (Omicron) for the EU/EEA. 26 Nov 2021. https://www.ecdc.europa.eu/sites/default/files/documents/Implications-emergence-spread-SARS-CoV-2%20B.1.1.529-variant-concern-Omicron-for-the-EU-EEA-Nov2021.pdf.

7. Siegel D, Hui HC, Doerffler E, Clarke MO, Chun K, Zhang L, Neville S, Carra E, Lew W, Ross B, Wang Q, Wolfe L, Jordan R, Soloveva V, Knox J, Perry J, Perron M, Stray KM, Barauskas O, Feng JY, Xu Y, Lee G, Rheingold AL, Ray AS, Bannister R, Strickley R, Swaminathan S, Lee WA, Bavari S, Cihlar T, Lo MK, Warren TK, Mackman RL. 2017. Discovery and synthesis of a phosphoramidate prodrug of a pyrrolo[2,1-f][triazin-4-amino] adenine C-nucleoside (GS-5734) for the treatment of Ebola and emerging viruses. J Med Chem 60:1648–1661.

8. Beigel JH, Tomashek KM, Dodd LE, Mehta AK, Zingman BS, Kalil AC, Hohmann E, Chu HY, Luetkemeyer A, Kline S, Lopez de Castilla D, Finberg RW, Dierberg K, Tapson V, Hsieh L Patterson TF, Paredes R, Sweeney DA, Short WR, Touloumi G, Lye DC, Ohmagari N, Oh MD, Ruiz-Palacios GM, Benfield T, Fatkenheuer G, Kortepeter MG, Atmar RL, Creech CB, Lundgren J, Babiker AG, Pett S, Neaton JD, Burgess TH, Bonnett T, Green M, Makowski M, Osinusi A, Nayak S, Lane HC, ACTT-1 Study Group Members. 2020. Remdesivir for the treatment of Covid-19 - Final report. N Engl J Med 383:1813–1826.

9. Goldman JD, Lye DCB, Hui DS, Marks KM, Bruno R, Montejano R, Spinner CD, Galli M, Ahn MY, Nahass RG, Chen YS, SenGupta D, Hyland RH, Osinusi AO, Cao H, Blair C, Wei X, Gaggar A, Brainard DM, Towner WJ, Munoz J, Mullane KM, Marty FM, Tashima KT, Diaz G, Subramanian A, GS-US-540-5773 Investigators. 2020. Remdesivir for 5 or 10 days in patients with severe Covid-19. N Engl J Med 383:1827–1837.

10. Spinner CD, Gottlieb RL, Criner GJ, Arribas Lopez JR, Cattelan AM, Soriano Viladomiu A, Ogbuagu O, Malhotra P, Mullane KM, Castagna A, Chai LYA, Roestenberg M, Tsang OTY, Bernasconi E, Le Turnier P, Chang SC, SenGupta D, Hyland RH, Osinusi AO, Cao H, Blair C, Wang H, Gaggar A, Brainard DM, McPhail MJ, Bhagani S, Ahn MY, Sanyal AJ, Huhn G, Marty FM, GS-US-540-5774 Investigators. 2020. Effect of remdesivir vs standard care on clinical status at 11 days in patients with moderate COVID-19: A randomized clinical trial. JAMA 324:1048–1057.

11. Gottlieb RL, Vaca CE, Paredes R, Mera J, Webb BJ, Perez G, Oguchi G, Ryan P, Nielsen BU, Brown M, Hidalgo A, Sachdeva Y, Mittal S, Osiyemi O, Skarbinski J, Juneja K, Hyland RH, Osinusi A, Chen S, Camus G, Abdelghany M, Davies S, Behenna-Renton N, Duff F, Marty FM, Katz MJ, Ginde AA, Brown SM, Schiffer JT, Hill JA, GS-US-540-9012 (PINETREE) Investigators. 2022. Early remdesivir to prevent progression to severe Covid-19 in outpatients. N Engl J Med 386:305–315.

12. US Food and Drug Administration (FDA). 2022. FDA News Release: FDA takes actions to expand use of treatment for outpatients with mild-to-moderate COVID-19. https://www.fda.gov/news-events/press-announcements/fda-takes-actions-expand-use-treatment-outpatients-mild-moderate-covid-19.

13. Pruijssers AJ, George AS, Schafer A, Leist SR, Gralinksi LE, Dinnon KH 3rd, Yount BL, Agostini ML, Stevens LJ, Chappell JD, Lu X, Hughes TM, Gully K, Martinez DR, Brown AJ, Graham RL, Perry JK, Du Pont V, Pitts J, Ma B, Babusis D, Murakami E, Feng JY, Bilello JP, Porter DP, Cihlar T, Baric RS, Denison MR, Sheahan TP. 2020. Remdesivir inhibits SARS-CoV-2 in human lung cells and chimeric SARS-CoV expressing the SARS-CoV-2 RNA polymerase in mice. Cell Rep 32:107940.

14. Gordon CJ, Tchesnokov EP, Woolner E, Perry JK, Feng JY, Porter DP, Gotte M. 2020. Remdesivir is a direct-acting antiviral that inhibits RNA-dependent RNA polymerase from severe acute respiratory syndrome coronavirus 2 with high potency. J Biol Chem 295:6785–6797.

15. Tchesnokov EP, Gordon CJ, Woolner E, Kocinkova D, Perry JK, Feng JY, Porter DP, Gotte M. 2020. Template-dependent inhibition of coronavirus RNA-dependent RNA polymerase by remdesivir reveals a second mechanism of action. J Biol Chem 295:16156–16165.

16. Cho A, Saunders OL, Butler T, Zhang L, Xu J, Vela JE, Feng JY, Ray AS, Kim CU. 2012. Synthesis and antiviral activity of a series of 1’-substituted 4-aza-7,9-dideazaadenosine C-nucleosides. Bioorg Med Chem Lett 22:2705–2707.

17. Warren TK, Jordan R, Lo MK, Ray AS, Mackman RL, Soloveva V, Siegel D, Perron M, Bannister R, Hui HC, Larson N, Strickley R, Wells J, Stuthman KS, Van Tongeren SA, Garza NL, Donnelly G, Shurtleff AC, Retterer CJ, Gharaibeh D, Zamani R, Kenny T, Eaton BP, Grimes E, Welch LS, Gomba L, Wilhelmsen CL, Nichols DK, Nuss JE, Nagle ER, Kugelman JR, Palacios G, Doerffler E, Neville S, Carra E, Clarke MO, Zhang L, Lew W, Ross B, Wang Q, Chun K, Wolfe L, Babusis D, Park Y, Stray KM, Trancheva I, Feng JY, Barauskas O, Xu Y, Wong P, Braun MR, Flint M, McMullan LK, Chen SS, Fearns R, Swaminathan S, Mayers DL, Spiropoulou CF, Lee WA, Nichol ST, Cihlar T, Bavari S. 2016. Therapeutic efficacy of the small molecule GS-5734 against Ebola virus in rhesus monkeys. Nature 531:381–385.

18. Mackman RL, Hu HC, Perron M, Murakami E, Palmiotti C, Lee G, Stray K, Zhang L, Goyal B, Chun K, Byun D, Siegel D, Simonovich S, Du Pont V, Pitts J, Babusis D, Vijjapurapu A, Lu X, Kim C, Zhao X, Chan J, Ma B, Lye D, Vandersteen A, Wortman S, Barrett KT, Toteva M, Jordan R, Subramanian R, Bilello JP, Cihlar T. 2021. Prodrugs of a 1’-CN-4-Aza-7,9-dideazaadenosine C-nucleoside leading to the discovery of remdesivir (GS-5734) as a potent inhibitor of respiratory syncytial virus with efficacy in the African green monkey model of RSV. J Med Chem 64:5001–5017.

19. Agostini ML, Andres EL, Sims AC, Graham RL, Sheahan TP, Lu X, Smith EC, Case JB, Feng JY, Jordan R, Ray AS, Cihlar T, Siegel D, Mackman RL, Clarke MO, Baric RS, Denison MR. 2018. Coronavirus susceptibility to the antiviral remdesivir (GS-5734) is mediated by the viral polymerase and the proofreading exoribonuclease. mBio 9:e00221–e00218.

20. Sheahan TP, Sims AC, Graham RL, Menachery VD, Gralinski LE, Case JB, Leist SR, Pyrc K, Feng JY, Trantcheva I, Bannister R, Park Y, Babusis D, Clarke MO, Mackman RL, Spahn JE, Palmiotti CA, Siegel D, Ray AS, Cihlar T, Jordan R, Denison MR, Baric RS. 2017. Broad-spectrum antiviral GS-5734 inhibits both epidemic and zoonotic coronaviruses. Sci Transl Med 9:eaa13653.

21. Sheahan TP, Sims AC, Zhou S, Graham RL, Pruijssers AJ, Agostini ML, Leist SR, Schafer A, Dinnon KH 3rd, Stevens LJ, Chappell JD, Lu X, Hughes TM, George AS, Hill CS, Montgomery, SA, Brown AJ, Bluemling GR, Natchus MG, Saindane M, Kolykhalov AA, Painter G, Harcourt J, Tamin A, Thornburg NJ, Swanstrom R, Denison MR, Baric RS. 2020. An orally bioavailable broad-spectrum antiviral inhibits SARS-CoV-2 in human airway epithelial cell cultures and multiple coronaviruses in mice. Sci Transl Med 12:eabb5883.

22. de Wit E, Feldmann F, Cronin J, Jordan R, Okumura A, Thomas T, Scott D, Cihlar T, Feldmann H. 2020. Prophylactic and therapeutic remdesivir (GS-5734) treatment in the rhesus macaque model of MERS-CoV infection. Proc Natl Acad Sci USA 117:6771–6776.

23. Lo MK, Feldmann F, Gary JM, Jordan R, Bannister R, Cronin J, Patel NR, Klena JD, Nichol ST, Cihlar T, Zaki SR, Feldmann H, Spiropoulou CF, de Wit E. 2019. Remdesivir (GS-5734) protects African green monkeys from Nipah virus challenge. Sci Transl Med 11:eaau9242.

24 Xie X, Muruato AE, Zhang X, Lokugamage KG, Fontes-Garfias CR, Zou J, Liu J, Ren P, Balakrishnan M, Cihlar T, Tseng CK, Makino S, Menachery VD, Bilello JP, Shi PY. 2020. A nanoluciferase SARS-CoV-2 for rapid neutralization testing and screening of anti-infective drugs for COVID-19. Nat Commun 11:5214.

25. Dai W, Zhang B, Jiang XM, Su H, Li J, Zhao Y, Xie X, Jin Z, Peng J, Liu F, Li C, Li Y, Bai F, Wang H, Chen X, Cen X, Hu S, Yang X, Wang J, Liu X, Xiao G, Jiang H, Rao Z, Zhang LK, Xu Y, Yang H, Liu H. 2020. Structure-based design of antiviral drug candidates targeting the SARS-CoV-2 main protease. Science 368:1331–1335.

26. Do TND, Donckers K, Vangeel L, Chatterjee AK, Gallay PA, Bobardt MD, Bilello JP, Cihlar T, De Jonghe S, Neyts J, Jochmans D. 2021. A robust SARS-CoV-2 replication model in primary human epithelial cells at the air liquid interface to assess antiviral agents. Antiviral Res 192:105122.

27. Li Y, Cao L, Li G, Cong F, Li Y, Sun J, Luo Y, Chen G, Li G, Wang P, Xing F, Ji Y, Zhao J, Zhang Y, Guo D, Zhang X. 2021. Remdesivir metabolite GS-441524 effectively inhibits SARS-CoV-2 infection in Mouse models. J Med Chem Feb 1:acs.jmedchem.0c01929.

28. Ye ZW, Yuan S, Chan JFW, Zhang AJ, Yu CY, Ong CP. 2021. Beneficial effect of combinational methylprednisolone and remdesivir in hamster model of SARS-CoV-2 infection. Emerg Microbes Infect 10:291–304.

29. Martinez DR, Schäfer A, Leist SR, Li D, Gully K, Yount B, Feng JY, Bunyan E, Porter DP, Cihlar T, Montgomery SA, Haynes BF, Baric RS, Nussenzweig MC, Sheahan TP. 2021. Prevention and therapy of SARS-CoV-2 and the B.1.351 variant in mice. Cell Rep 36:109450.

30. Williamson BN, Feldmann F, Schwarz B, Meade-White K, Porter DP, Schulz J, van Doremalen N, Leighton I, Yinda CK, Pérez-Pérez L, Okumura A, Lovaglio J, Hanley PW, Saturday G, Bosio CM, Anzick S, Barbian K, Cihlar T, Martens C, Scott DP, Munster VJ, de Wit E. 2020. Clinical benefit of remdesivir in rhesus macaques infected with SARS-CoV-2. Nature 585:273–276.

31. Vermillion MS, Murakami E, Ma B, Pitts J, Tomkinson A, Rautiola D, Babusis D, Irshad H, Siegel D, Kim C, Zhao X, Niu C, Yang J, Gigliotti A, Kadrichu N, Bilello JP, Ellis S, Bannister R, Subramanian R, Smith B, Mackman RL, Lee WA, Kuehl PJ, Hartke J, Tomas Cihlar T, Porter DP. 2021. Inhaled remdesivir reduces viral burden in a nonhuman primate model of SARS-CoV-2 infection. Sci Transl Med 10.1126/scitranslmed.ab18282

32. Williamson BN, Pérez-Pérez L, Schwarz B, Feldmann F, Holbrook MG, Singh M, Lye DS, Babusis D, Subramanian R, Haddock E, Okumura A, Hanley PW, Lovaglio J, Bosio CM, Porter DP, Cihlar T, Mackman RL, Saturday G, de Wit E. 2022. Subcutaneous remdesivir administration prevents interstitial pneumonia in rhesus macaques inoculated with SARS-CoV-2. Antiviral Res 198:105246.

33. Martin R, Li J, Parvangada A, Perry J, Cihlar T, Mo H, Porter D, Svarovskaia E. 2021. Genetic conservation of SARS-CoV-2 RNA replication complex in globally circulating isolates and recently emerged variants from humans and minks suggests minimal pre-existing resistance to remdesivir. Antivir Res 188:105033.

34. Zhao H, Lu L, Peng Z, Chen LL, Meng X, Zhang C, Ip JD, Chan WM, Chu AW, Chan KH, Jin DY, Chen H, Yuen KY, To KK. 2022. SARS-CoV-2 Omicron variant shows less efficient replication and fusion activity when compared with Delta variant in TMPRSS2-expressed cells. Emerg Microbes Infect 11:277–283.

35. Szemiel AM, Merits A, Orton RJ, MacLean OA, Pinto RM, Wickenhagen A, Lieber G, Turnbull ML, Wang S, Furnon W, Suarez NM, Mair D, da Silva Filipe A, Willett BJ, Wilson SJ, Patel AH, Thomson EC, Palmarini M, Kohl A, Stewart ME. 2021. In vitro selection of remdesivir resistance suggests evolutionary predictability of SARS-CoV-2. PLoS Pathog 17:e1009929.

36. Gordon, CJ, Lee, HW, Tchesnokov, EP, Perry, JK, Feng, JY, Bilello, JP, Porter, DP, Götte, M. 2022. Efficient incorporation and template-dependent polymerase inhibition are major determinants for the broad-spectrum antiviral activity of remdesivir. J Biol Chem 298:101529.

37. Chen J, Malone B, Llewellyn E, Grasso M, Shelton PMM, Olinares PDB, Maruthi K, Eng ET, Vatandaslar H, Chait BT, Kapoor TM, Darst SA, Campbell EA. 2020. Structural basis for helicase-polymerase coupling in the SARS-CoV-2 replication-transcription complex. Cell 182:1560–1573.e13.

38. Beard H, Cholleti A, Pearlman D, Sherman W, Loving KA. 2013. Applying physics-based scoring to calculate free energies of binding for single amino acid mutations in protein-protein complexes. PLoS One 8:e82849.

39. Dong X, Goldswain H, Penrice-Randal R, Shawli GT, Prince T, Kavanagh Williamson M, Randle N, Jones B, Salguero FJ, Tree JA, Hall Y, Hartley C, Erdmann M, Bazire J, Jearanaiwitayakul T, ISARIC4C Investigators, Semple MG, Openshaw PJM, Baille JK, Emmett SR, Digard P, Matthews DA, Turtle L, Darby A, Davidson AD, Carroll MW, Hiscox JA. 2021. Rapid selection of P323L in the SARS-CoV-2 polymerase (NSP12) in humans and non-human primate models and confers a large plaque phenotype. bioRxiv 2021.12.23.474030.

40. Cox RM, Wolf JD, Lieber CM, Sourimant J, Lin MJ, Babusis D, DuPont V, Chan J, Barrett KT, Lye D, Kalla R, Chun K, Mackman RL, Ye C, Cihlar T, Martinez-Sobrido L, Greninger AL, Bilello JP, Plemper RK. 2021. Oral prodrug of remdesivir parent GS-441524 is efficacious against SARS-CoV-2 in ferrets. Nat Commun 12:6415.

41. Schäfer A, Martinez DR, Won JJ, Moreira FR, Brown AJ, Gully KL, Kalla R, Chun K, Du Pont V, Babusis D, Tang J, Murakami E, Subramanian R, Barrett KT, Bleier BJ, Bannister R, Feng JY, Bilello JP, Cihlar T, Mackman RL, Montgomery SA, Baric RS, Sheahan TP. 2021. Therapeutic efficacy of an oral nucleoside analog of remdesivir against SARS-CoV-2 pathogenesis in mice. bioRxiv 2021.09.13.460111.

42. Focosi D, Maggi F, McConnell S, Casadevall A. 2022. Very low levels of remdesivir resistance in SARS-COV-2 genomes after 18 months of massive usage during the COVID19 pandemic: A GISAID exploratory analysis. Antiviral Res 198:105247.

43. Cao Y, Wang J, Jian F, Xiao T, Song W, Yisimayi A, Huang W, Li Q, Wang P, An R, Wang J, Wang Y, Niu X, Yang S, Liang H, Sun H, Li T, Yu Y, Cui Q, Liu S, Yang X, Du S, Zhang Z, Hao X, Shao F, Jin R, Wang X, Xiao J, Wang Y, Xie XS. 2021. Omicron escapes the majority of existing SARS-CoV-2 neutralizing antibodies. Nature 10.1038/s41586-021-04385-3.

44. Muik A, Lui BG, Wallisch AK, Bacher M, Mühl J, Reinholz J, Ozhelvaci O, Beckmann N, Güimil Garcia RC, Poran A, Shpyro S, Finlayson A, Cai H, Yang Q, Swanson KA, Türeci Ö, Şahin U. 2022. Neutralization of SARS-CoV-2 Omicron by BNT162b2 mRNA vaccine-elicited human sera. Science eabn7591.

45. Shirogane Y, Takeda M, Iwasaki M, Ishiguro N, Takeuchi H, Nakatsu Y, Tahara M, Kikuta H, Yanagi Y. 2008. Efficient multiplication of human metapneumovirus in Vero cells expressing the transmembrane serine protease TMPRSS2. J Virol 82:8942–8946.

46. Mossel EC, Huang C, Narayanan K, Makino S, Tesh RB, Peters CJ. 2005. Exogenous ACE2 expression allows refractory cell lines to support severe acute respiratory syndrome coronavirus replication. J Virol 79:3846–3850.

47. Elbe S, Buckland-Merrett G. 2017. Data, disease and diplomacy: GISAID’s innovative contribution to global health. Glob Chall 1:33–46.

48. O’Toole Á, Scher E, Underwood A, Jackson B, Hill V, McCrone JT, Colquhoun R, Ruis C, Abu-Dahab K, Taylor B, Yeats C, du Plessis L, Maloney D, Medd N, Attwood SW, Aanensen DM, Holmes EC, Pybus OG, Rambaut A. 2021. Assignment of epidemiological lineages in an emerging pandemic using the pangolin tool. Virus Evol 7:veab064.

49. Xie X, Lokugamage KG, Zhang X, Vu MN, Muruato AE, Menachery VD, Shi PY. 2021. Engineering SARS-CoV-2 using a reverse genetic system. Nat Protoc 16:1761–1784.

50. Xie X, Muruato A, Lokugamage KG, Narayanan K, Zhang X, Zou J, Liu J, Schindewolf C, Bopp NE, Aguilar PV, Plante KS, Weaver SC, Makino S, LeDuc JW, Menachery VD, Shi PY. 2020. An Infectious cDNA Clone of SARS-CoV-2. Cell Host Microbe 27:841–848.e3.

51. Liu Y, Liu J, Johnson BA, Xia H, Ku Z, Schindewolf C, Widen SG, An Z, Weaver SC, Menachery VD, Xie X, Shi PY. 2021. Delta spike P681R mutation enhances SARS-CoV-2 fitness over Alpha variant. bioRxiv 2021.08.12.456173.

